# Visual Cell Sorting: A High-throughput, Microscope-based Method to Dissect Cellular Heterogeneity

**DOI:** 10.1101/856476

**Authors:** Nicholas Hasle, Anthony Cooke, Sanjay Srivatsan, Heather Huang, Jason J. Stephany, Zachary Krieger, Dana Jackson, Weiliang Tang, Sriram Pendyala, Raymond J. Monnat, Cole Trapnell, Emily M. Hatch, Douglas M. Fowler

## Abstract

Microscopy is a powerful tool for characterizing complex cellular phenotypes, but linking these phenotypes to genotype or RNA expression at scale remains challenging. Here, we present Visual Cell Sorting, a method that physically separates hundreds of thousands of live cells based on their visual phenotype. Visual Cell Sorting uses automated imaging and phenotypic analysis to direct selective illumination of Dendra2, a photoconvertible fluorescent protein expressed in live cells; these photoactivated cells are then isolated using fluorescence-activated cell sorting. First, we use Visual Cell Sorting to assess the effect of hundreds of nuclear localization sequence variants in a pooled format, identifying variants that improve nuclear localization and enabling annotation of nuclear localization sequences in thousands of human proteins. Second, we use Visual Cell Sorting to recover cells that retain normal nuclear morphologies after paclitaxel treatment, then derive their single cell transcriptomes to identify multiple pathways associated with paclitaxel resistance in human cancers. Unlike alternative methods, Visual Cell Sorting depends on inexpensive reagents and commercially available hardware. As such, it can be readily deployed to uncover the relationships between visual cellular phenotypes and internal states, including genotypes and gene expression programs.

## Introduction

High content imaging^1^, *in situ* sequencing methods^2–9^, and other approaches^10–14^ have revolutionized the investigation of how genetic variants and gene expression programs dictate cellular morphology, organization and behavior. One important application of these methods is visual genetic screening, in which a library of genetic variants is introduced into cells and the effect of each variant on a visual phenotype is quantified. In a classical high content visual genetic screen, each genetic perturbation occupies a separate well. New *in situ* methods, which employ sequencing by repeated hybridization of fluorescent oligo probes^2–6^ or direct synthesis^7–9, 15^ to visually read out nucleic acid barcodes, permit hundreds of perturbations to be assessed in a pooled format. For example, multiplexed fluorescent in-situ hybridization was used to assess the effect of 210 CRISPR sgRNAs on RNA localization in ∼30,000 cultured human U-2 OS cells^5^; and *in situ* sequencing was used to measure the effect of 963 gene knockouts on the localization of an NFkB reporter at a throughput of ∼3 million cells^16^. Visual phenotyping methods can also dissect non-genetic drivers of phenotypic heterogeneity. Here, characterization of cells with distinct visual phenotypes can reveal different cell states – such as signaling pathway activities and gene expression profiles – that are associated with different cellular morphologies. For example, the photoactivatable marker technology Single-Cell Magneto-Optical Capture was used to isolate and sequence the transcriptomes of cells that successfully resolved ionizing radiation-induced DNA damage foci^12^.

Despite their utility, current methods have limitations (Supplementary Table 1). Some, such as high content imaging require highly specialized or custom-built hardware. Others, like *in situ* sequencing, employ complex protocols, sophisticated computational pipelines, and expensive dye-based reagents. Methods that mark and sort for individual cells with a photoactivatable protein or compound are simpler and less expensive. However, are either low throughput (< 1,000 cells per experiment)^10–13^ or lack single-cell specificity^14^. Furthermore, they cannot investigate more than one or two phenotypes per experiment.

To address these shortcomings, we developed Visual Cell Sorting, a flexible and simple high-throughput method that uses commercial hardware to enable the investigation of cells according to visual phenotype. Visual Cell Sorting is an automated platform that directs a digital micromirror device to mark single live cells that express a nuclear photoactivatable fluorescent protein for subsequent physical separation by fluorescence activated cell sorting (FACS). We demonstrate that Visual Cell Sorting enables visual phenotypic sorting into 4 bins; increases the throughput of cellular separation by 1,000-fold compared to other single cell photoconversion-based technologies^10–13^; and permits pooled genetic screening and transcriptomic profiling. For example, Visual Cell Sorting enabled us to sort hundreds of thousands of cultured human cells according to the nuclear localization of a fluorescent reporter protein, and thus score a library of nuclear localization sequence variants for function. In a second application, we isolated paclitaxel-treated cells with normal or lobulated nuclear morphologies and subjected each population to single cell RNA sequencing, revealing multiple pathways associated with paclitaxel resistance. Visual Cell Sorting requires simple, inexpensive, and commercially available widefield microscope hardware, routine genetic engineering, and a standard 4-laser FACS instrument to perform. As such, we envision that Visual Cell Sorting can readily be deployed to uncover the relationships between visual cellular phenotypes and their associated internal states, including genotype and gene expression programs.

## Results

### Physical Separation of Cells by Visual Phenotype

Visual Cell Sorting uses FACS to separate hundreds of thousands of cells by their visual phenotypes. Cells are first modified to express Dendra2, a green-to-red photoconvertible fluorescent protein^17^ that will act as a phenotypic marker and enable downstream FACS sorting. Next, cells are imaged on an automated microscope. In each field of view, cells are identified and analyzed for phenotypes of interest. According to their phenotype, cells are illuminated with 405 nm light for different lengths of time using a digital micromirror device, resulting in different levels of red Dendra2 fluorescence. The imaging, analysis, and photoactivation steps are performed at each field of view; and unlike previous photoactivatable marker-based methods, these steps are automated, allowing hundreds of thousands of cells to be assessed per experiment. Once all cells have been imaged, analyzed, and photoactivated, FACS is used to sort them into bins according to their level of Dendra2 photoactivation (Figure 1A). We first sought to establish the single cell accuracy of Dendra2 photoactivation, and whether variable photoactivation states could be discerned by flow cytometry. We noticed that similar technologies use photoactivatable dyes or proteins localized to the whole cell body^10–13^. This localization strategy makes identifying the boundaries of the fluorescent signal difficult, which results in partial photoactivation or photoactivation of the marker in a cell adjacent to a cell of interest. With this in mind, we expressed Dendra2 in the nucleus either as a histone H3 fusion (H3-Dendra2) or with an upstream nuclear localization sequence (NLS-Dendra2×3). The boundaries of nuclear Dendra2 signal are easy to identify, permitting quantitative photoactivation of Dendra2 in the cells of interest; and the cytoplasm provides a spacer between the Dendra2 in different cells, reducing photoactivation of cells adjacent to the cells of interest.

**Figure 1.**
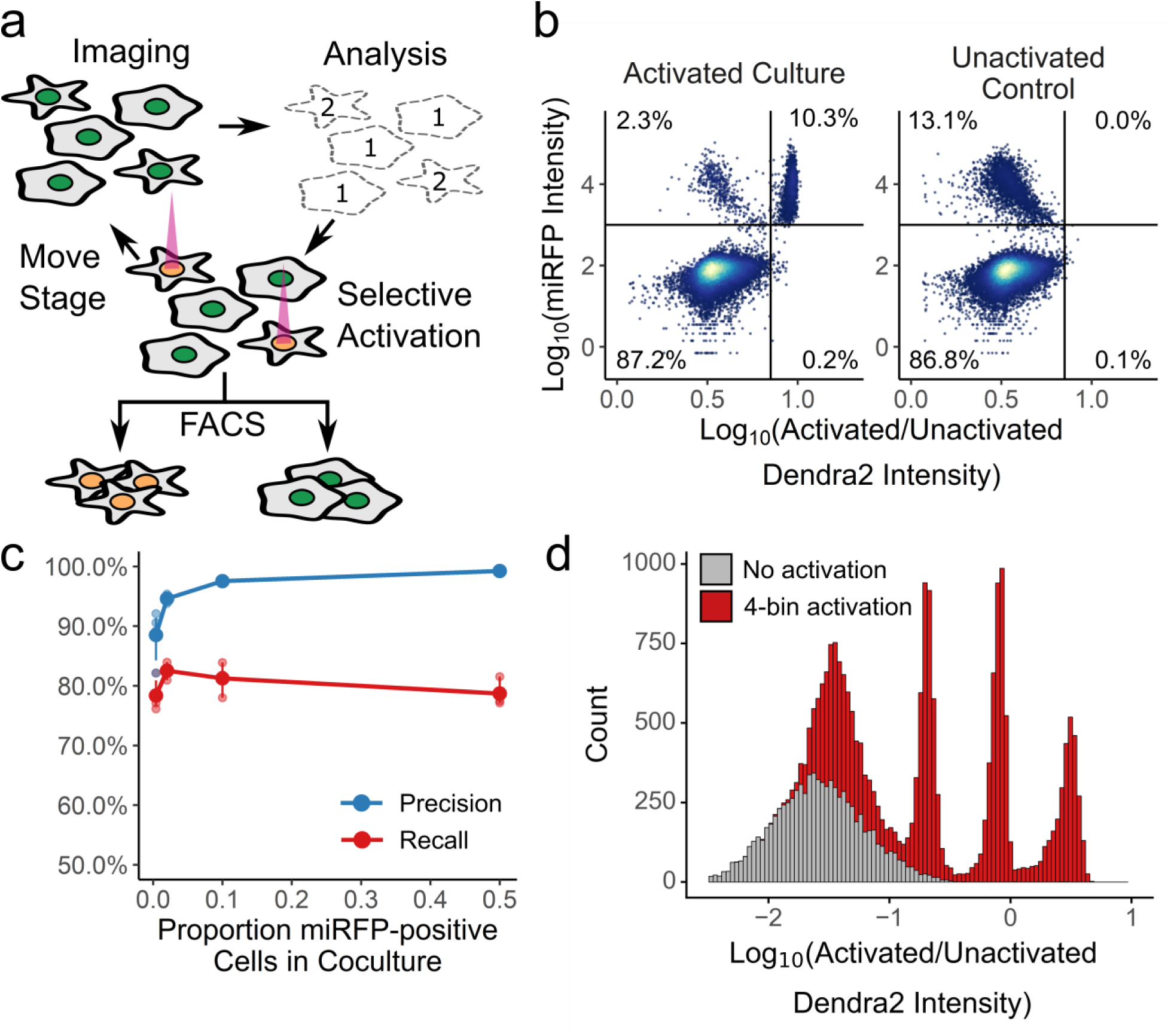
Visual Cell Sorting. **(a)** In an automated fashion, cells in a field of view are imaged and their phenotype classified. Cells of interest are illuminated with 405 nm light, which irreversibly photoactivates Dendra2 from its green to its red fluorescent state. The microscope then moves to a new field of view. These steps are repeated across an entire culture well. Then, fluorescence-activated cell sorting based on Dendra2 photoactivation is used to physically recover cells of interest. **(b)** To assess the photoactivation accuracy, U-2 OS cells expressing nuclear Dendra2 and miRFP, or nuclear Dendra2 alone, were co-cultured. The microscope was programmed to activate Dendra2 in cells expressing miRFP. Following photoactivation, miRFP expression and the ratio of activated to unactivated Dendra2 (left panel, n = 18,766 cells) were assessed with flow cytometry. In a second co-culture, Dendra2 was unactivated (right panel, n = 18,395 cells). Lines indicate gates for miRFP-positive cells and activated Dendra2 cells, with the percentage of cells appearing in each quadrant indicated. **(c)** Same experiment as (b), except cells were mixed such that 0.5%, 4%, 12%, or 50% were miRFP positive. Precision and recall were computed; large solid, mean (N = 3 replicates); small points, individual replicate values; error bars, standard error from the mean. **(d)** U-2 OS cells in one well were illuminated with 405 nm light for 0, 50, 200, or 800 ms (red; n = 16,397). Cells in a second well were left unactivated (grey; n = 8,497). The ratio of activated to unactivated Dendra2 was determined by flow cytometry.

To measure photoactivation accuracy, H3-Dendra2 positive cells co-expressing H2B-miRFP were mixed with cells expressing H3-Dendra2 alone at decreasing ratios. We instructed the microscope to activate Dendra2 in cells harboring miRFP-positive nuclei, and then we quantified the co-occurrence of miRFP and activated Dendra2 florescence signals using flow cytometry (Figure 1B). The ratio of activated Dendra2 fluorescence to unactivated Dendra2 fluorescence (Dendra2 photoactivation ratio) accurately predicted whether a cell was miRFP-positive, even when the miRFP expressing cells were present at ∼0.5% frequency, with average precision of 94% and recall of 80% (Figure 1C).

Previous photoactivatable marker-based methods have been limited to two photactivation levels: activated and unactivated. To test whether we could encode more than one photoactivation level, and thus more than one phenotype, we exposed different cells in the same well to 405 nm light for 0, 50, 200, or 800 ms. Flow cytometry of the Dendra2 fluorescence distribution by showed four distinct levels of Dendra2 photoactivation, indicating that Visual Cell Sorting can sort four different visual phenotypes or four discrete bins of a continuous phenotype (Figure 1D). Furthermore, these four photoactivation levels can still be distinguished over 12 hours following activation (Supplementary Figure 1). To extend the amount of time that the photoactivation levels remain distinct from one another, we placed H3-Dendra2 expression under the control of a doxycycline-inducible promoter. By shutting off Dendra2 expression before the experiment, the 50, 200, and 800 ms photoactivation levels remained distinguishable for up to 24 hours (Supplementary Figure 1). Finally, we examined the effect of Dendra2 photoactivation on cell viability and function. Activated cells did not exhibit higher rates of apoptosis or cell death even two days after photoactivation, nor did we detect effects of photoactivation on gene expression (Supplementary Figure 1). These results indicate that Dendra2 photoactivation does not appreciably affect cell survival or gene expression programs.

### Visual Cell Sorting Enables Pooled, Image-based Genetic Screening

To test whether Visual Cell Sorting enables image-based genetic screening, we asked if we could separate cells according to the nuclear localization of a fluorescent reporter protein. Nuclear localization sequences (NLS’s) are short peptides that direct proteins to the nucleus, and NLS’s are critical for the function of thousands of human transcription factors, nuclear structural proteins, and chromatin modifying enzymes. Over 90% of nuclear proteins do not have an annotated nuclear localization sequence in UniProt, and current NLS prediction algorithms cannot sensitively identify known NLS’s without drastically decreasing their precision^18, 19^. This shortcoming may arise because these NLS prediction algorithms rely on sequence alignments or amino acid frequencies of naturally observed NLS’s, which are subject to discovery bias. Therefore, we used Visual Cell Sorting to evaluate a large library of NLS missense variants; sort cells according to the NLS function; and sequence the sorted cells (Figure 2A), with the hypothesis that the resulting data could be used to improve NLS prediction.

**Figure 2.**
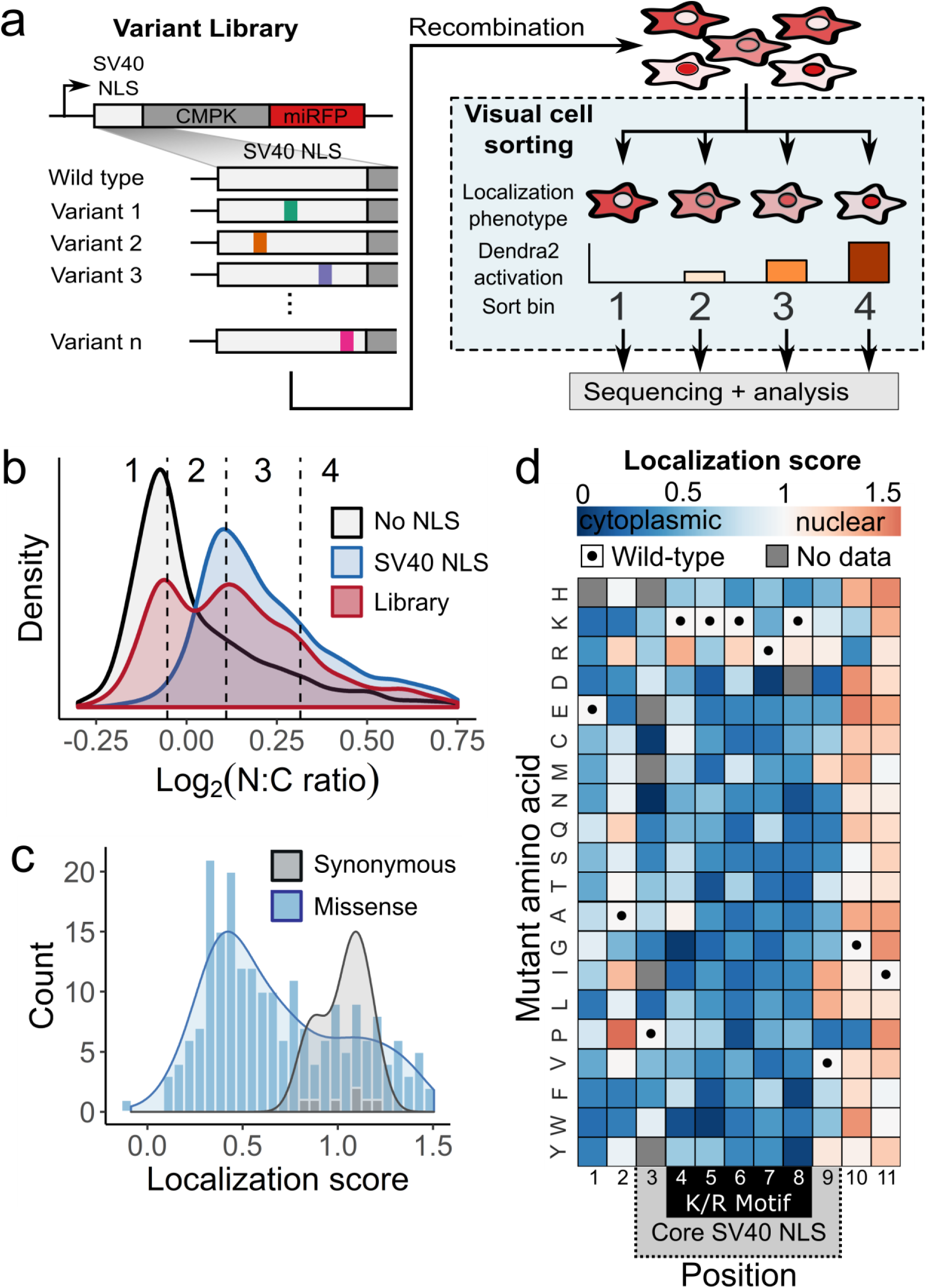
Visual Cell Sorting for pooled, image-based genetic screening. **(a)** A mutagenized simian virus (SV) 40 NLS library containing 346 unique nucleotide variants fused to a chicken muscle pyruvate kinase (CMPK) miRFP reporter was recombined into a U-2 OS H3-Dendra2 cell line. Visual Cell Sorting was performed to separate the NLS library expressing cells into four photoactivation bins according to the microscope-derived nucleus-to-cytoplasm ratio of the miRFP reporter. Each bin was deeply sequenced and analyzed to assign each amino acid variant a quantitative nuclear localization score. **(b)** U-2 OS H3-Dendra2 cells expressing either the NLS library, a wild type control or a no NLS control were imaged at 20X magnification and nucleus to cytoplasm (N:C) ratios measured. Curves, estimated kernel density of cells (n = 1,529 – 3,931 per condition); dotted lines, Visual Cell Sorting photoactivation gates with associated bin numbers. **(c)** Raw variant nuclear localization scores were calculated using a scaled weighted average of variant frequencies across the four sort bins. WT-like variants have a score of 1 and cytoplasm-localized variants a score of 0. Localization score, mean values of normalized scores from 5 replicates (N = 637,605 cells); curves, kernel density estimate of variant score distributions. **(d)** Nuclear localization scores of missense variants (N = 202) displayed as a heatmap. Gray boxes, variants not observed or scored in a single replicate; black dots, WT sequence; dotted gray area on the horizontal axis, SV40 NLS often used to localize recombinant proteins to the nucleus; black box, the five residue K/R-rich region.

We based our library on the SV40 NLS, a 7-residue sequence containing a lysine and arginine-rich region (K/R motif) that was the first NLS to be discovered^20^. To assess NLS variant function, we constructed a fluorescent nuclear localization reporter similar to one described previously^20^. Cultured U-2 OS H3-Dendra2 cells expressing the wild-type SV40 NLS fused to a CMPK-miRFP reporter had high levels of miRFP in the nucleus, relative to the cytoplasm. The degree of nuclear localization was calculated using a nucleus-to-cytoplasm miRFP intensity ratio (N:C ratio; Supplementary Figure 2). In contrast to the wild-type SV40 NLS-tagged reporter, cells expressing an untagged reporter had a low nucleus-to-cytoplasm ratio (Figure 2B).

We generated a library of 346 NLS nucleotide variants, corresponding to all possible 209 single amino acid missense variants. Cells expressing the library had a bimodal nucleus-to-cytoplasm ratio distribution, indicating that some variants preserved reporter nuclear localization while others disrupted its localization to different degrees (Figure 2B). We divided the library into four photoactivation levels spanning the nucleus-to-cytoplasm ratio range and used Visual Cell Sorting to sort cells into four bins (Supplementary Figure 2). A total of 637,605 cells were sorted across 5 replicates (Supplementary Table 2). Microscopy on the sorted cells revealed that Visual Cell Sorting faithfully separated cells by the nuclear localization phenotype (Supplementary Figure 2). Deep sequencing revealed the frequency of each variant in every bin, and we used these frequencies to compute a quantitative nuclear localization score for 97% of the 209 possible single missense variants (Supplementary Table 3)^21^. Scores were subsequently normalized such that wild-type had a normalized score of 1 and the bottom 10% of scoring variants had a median normalized score of 0.

As expected, nuclear localization scores for synonymous variants were close to a wild-type-like score of one, and most missense scores were lower than one, indicating loss of nuclear localization sequence function (Figure 2C). Furthermore, the SV40 NLS was most sensitive to substitutions in its K/R motif (Figure 2D). Localization scores were reproducible (mean r = 0.73; Supplementary Figure 2), and individually assessed nucleus-to-cytoplasm ratios were highly correlated to the localization scores derived using Visual Cell Sorting (r^2^ = 0.91; Figure 3A). Finally, localization scores of individual variants were correlated with previously reported *in vitro* K_d_ values for binding to importin alpha (r = −0.76, Supplementary Figure 3). Thus, Visual Cell Sorting accurately quantified the effect of NLS variants on their nuclear localization function.

**Figure 3.**
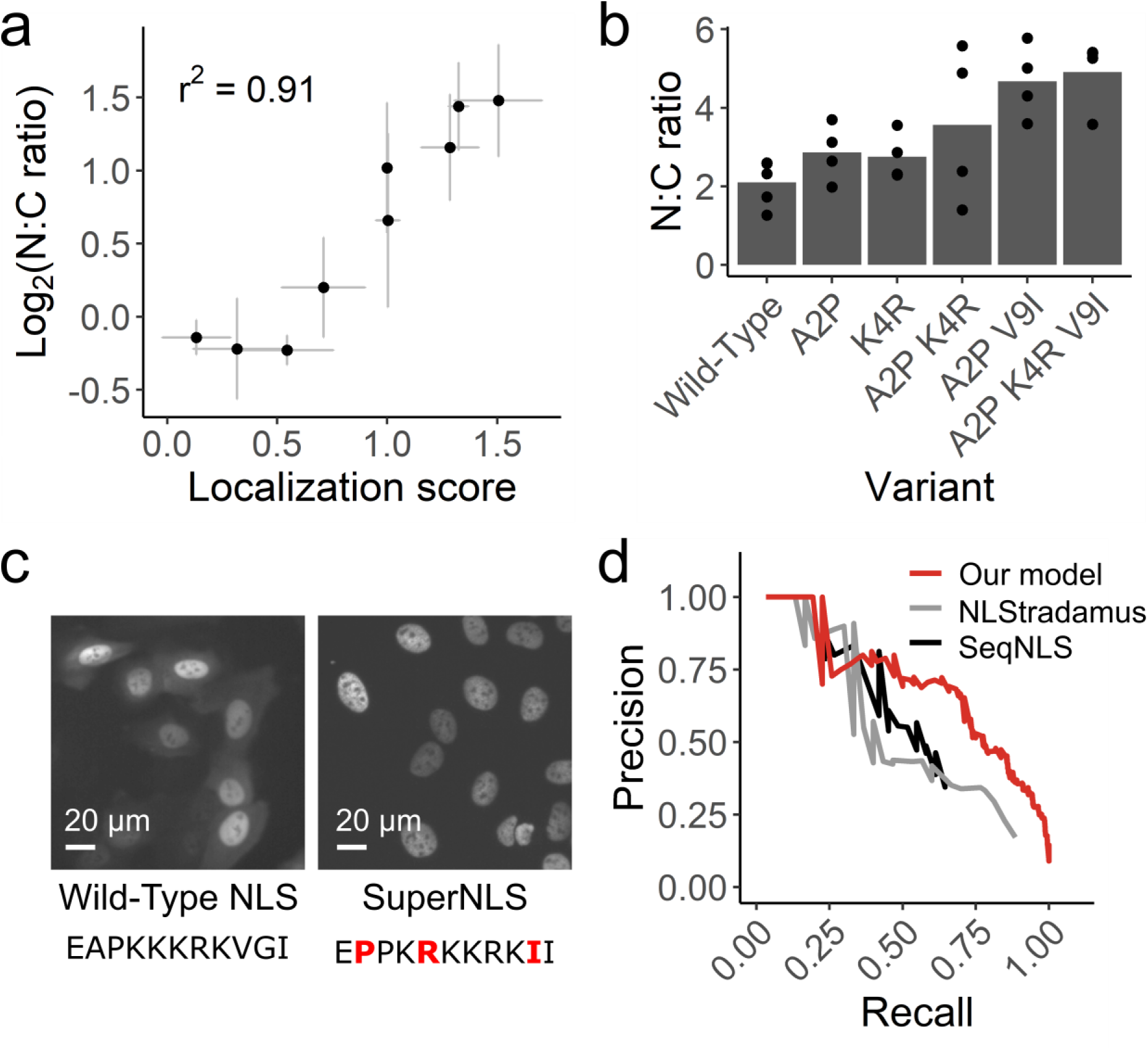
Visual Cell Sorting-derived variant scores accurately predict NLS function. **(a)** Nine NLS variants were individually expressed in the CMPK-miRFP reporter in U-2 OS H3-Dendra2 cells. The median nucleus to cytoplasm (N:C) ratio of cells expressing each variant was measured by microscope and compared to its localization score derived by Visual Cell Sorting. n ≥ 141 cells per variant per replicate. Bars, mean across at least three separate replicates. **(b)** SV40 NLS variants that appeared to enhance nuclear localization were individually tested both alone and in combination. NLS variants with up to three amino acid changes were expressed in U-2 OS H3-Dendra2 cells and imaged; the median N:C ratio was quantified across cells in the same well. n ≥ 527 cells per variant per replicate. **(c)** Representative images from cells expressing the wild-type SV40 NLS or the optimized superNLS fused to the miRFP reporter. Red letters, amino acid differences from wild-type. **(d)** Nuclear localization scores derived from Visual Cell Sorting were used to generate a predictive model that was trained on UniProt NLS annotations. Precision/recall curves for our model and two other linear motif scoring models, NLStradamus (Nguyen Ba et al. 2009) and SeqNLS (Lin et al. 2013), on a test dataset (n = 30 NLSs) are shown.

The SV40 NLS is commonly used to localize recombinant proteins to the nucleus and is included in over 10% of all constructs deposited in AddGene (accessed June 2019). Thus, an optimized NLS could improve a wide range of experiments including CRISPR-mediated genome editing. We further investigated three variants that appreciably increased nuclear localization of the reporter compared to the wild-type SV40 NLS. Individually, these variants modestly improved nuclear localization, and a “superNLS” with three missense variants increased nuclear localization by 2.3 fold (Figure 3B, C).

Most NLS prediction algorithms use naturally occurring, individually validated NLS sequences to identify similar sequences in new proteins. By contrast, our data comprise a comprehensive set of NLS-like sequences with variable function. We trained a linear regression model to predict whether any given 11-mer functions as a monopartite NLS by using the experimentally-determined amino acid preferences^22^ at each NLS position, which were calculated with the localization score data. We evaluated our model using a test dataset, not used for training, of 30 NLS’s in 20 proteins. The resulting model more accurately predicted NLS’s than two previously-published linear motif scoring models^18, 19^, particularly at a stringency where the majority of NLS’s are detected (Figure 3D). We used our model to annotate NLS’s in nuclear human proteins^23^ according to two score thresholds: one for high confidence monopartite NLS (precision 0.88, recall 0.23) and one for candidate monopartite NLS’s (precision 0.63, recall 0.70). In total, we annotated 2,310 high confidence monopartite NLS’s and an additional 20,013 candidate monopartite NLS’s across 6,714 human nuclear proteins (Supplementary Table 4).

### Visual Cell Sorting Enables Transcriptome Profiling on Image-based Phenotypes

To test whether Visual Cell Sorting enables transcriptomics on cells with distinct image-based phenotypes, we performed single cell RNA sequencing on cells undergoing divergent morphologic responses to paclitaxel. Paclitaxel is a chemotherapeutic agent that stabilizes microtubules and has been used to treat cancer for decades^24^. Even in a clonal population, a subset of cells adopt a lobulated nuclear morphology when treated with a low dose (≤ 10 nM) of paclitaxel^25^. We treated a telomerase-immortalized cell line derived from human retinal pigment epithelium, hTERT RPE-1, with paclitaxel and observed mitoses that sometimes resulted in nuclear lobulation that persists through the cell cycle (Videos S1, S2). In order to computationally define a cutoff for lobulated nuclei, we measured the shape factor, a circularity metric (Figure 4A), of nuclei in vehicle-treated cells and found that 95% of these morphologically normal cells have a nuclear shape factor greater than 0.65. We then analyzed paclitaxel treated cells and found that 30% of paclitaxel treated cells had lobulated nuclei, defined by shape factor of less than 0.65 (Figure 4B).

**Figure 4.**
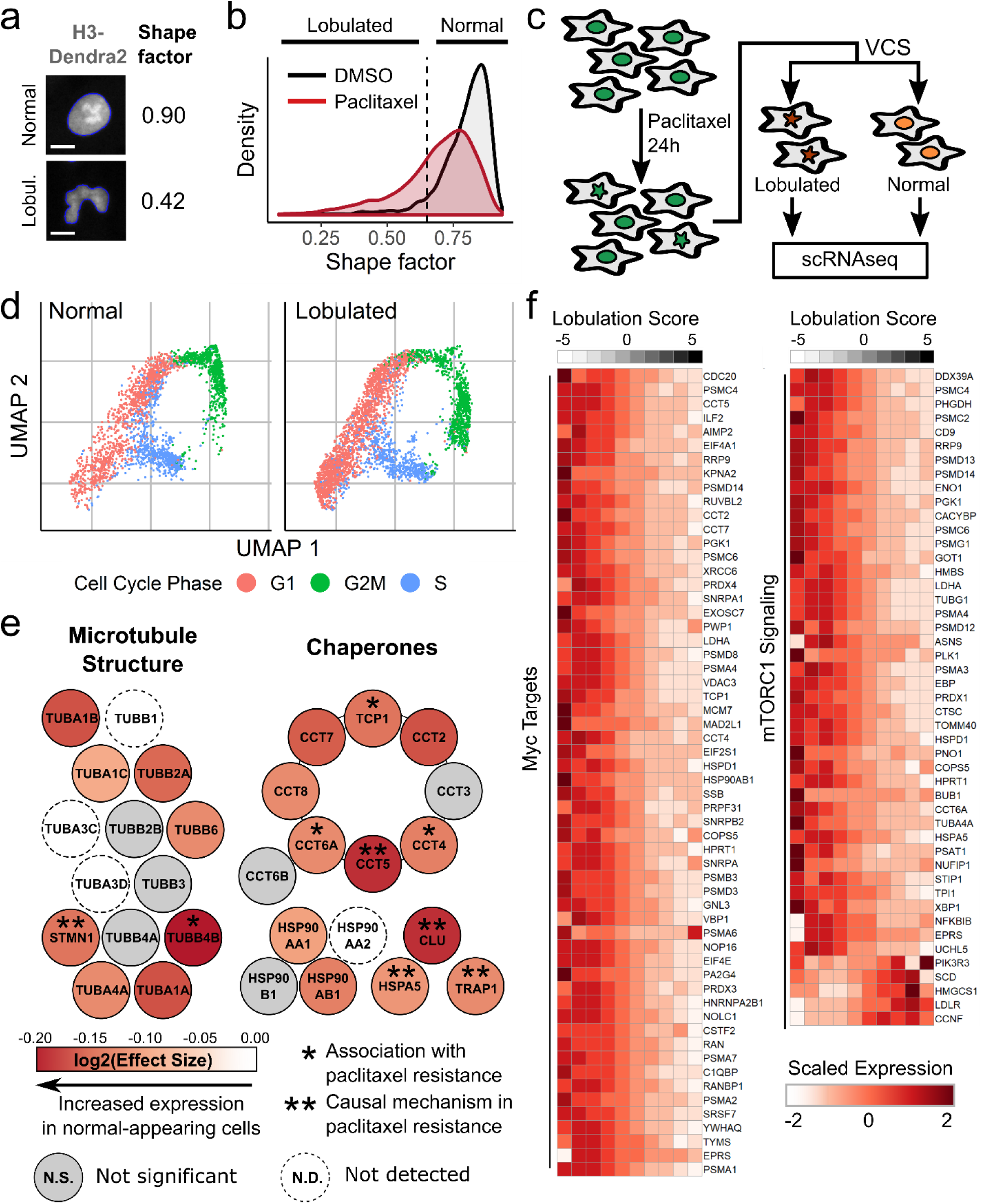
Visual Cell Sorting to dissect heterogeneous nuclear morphology following paclitaxel treatment. **(a)** RPE-1 NLS-Dendra2×3 cells were treated for 30 hours with 0.25 nM paclitaxel or DMSO and imaged. The shape factor, which measures the degree of an object’s circularity, was computed for each nucleus. One normal nucleus with a shape factor near one and one lobulated nucleus with a low shape factor are shown. The computationally determined boundaries of each nucleus are shown in blue; scale bar = 10 μm. **(b)** Shape factor density plots for vehicle (DMSO) and 0.25 nM paclitaxel-treated RPE-1 cells (n ≥ 3,914 cells per treatment). Dashed line, cutoff for lobulated nuclei (shape factor < 0.65). **(c)** RPE-1 cells were treated with 0.25 nM paclitaxel, then subjected to Visual Cell Sorting according to nuclear shape factor. Populations of cells with normal or lobulated nuclei were subjected separately to single cell RNA sequencing. **(d)** UMAP analysis of single cell RNA sequencing results. Expression of cell-cycle related genes were used to annotate each cell as being in G1, S, or G2/M. **(e)** A differential gene test was performed using as covariates cell cycle scores and a lobulation score, which is higher in lobulated cells compared to morphologically normal cells (Supplementary Figure 4). Genes related to microtubule structure or various chaperone complexes are colored according to the expected log2 fold-change per unit increase in lobulation score (effect size); asterisks, genes associated with paclitaxel resistance^34, 37, 39, 59–61^. **(f)** Expression counts for genes associated with c-Myc and mTORC1 signaling were aggregated across cells binned according to their lobulation score, then log-normalized and rescaled. Higher lobulation scores correspond to a higher likelihood of nuclear lobulation.

Given that morphologic phenotypes are potent indicators of cell state^26^, we hypothesized that the change in nuclear morphology was accompanied by a distinct gene expression program. To test this hypothesis, we used Visual Cell Sorting to separate morphologically normal paclitaxel-treated cells (shape factor > 0.65) from those with lobulated nuclei (shape factor < 0.65). We then subjected each population of cells to single cell RNA sequencing (Figure 4C). Imaging, analysis, photoactivation, and FACS-based recovery (Supplementary Figure 4) of ∼200,000 cells took less than 7 hours. Following FACS, we prepared sequencing libraries for approximately 6,000 single cell transcriptomes from each population. We observed an RNA sequencing batch effect that was completely attributable to different levels of cell-free RNA^27, 28^ in the lobulated and morphologically-normal cell sequencing preps (Supplementary Figure 4).

We used UMAP^29^ to visualize a low-dimensional embedding of the single cell transcriptomes. The distributions of normal and lobulated cells in the UMAP embedding were similar, indicating modest differences in their transcriptomic states. Differences in cell-cycle phase^30^ largely explained transcriptomic variation (Figure 4D). More lobulated cells than normal cells were in G1 (53% vs. 44%), suggesting that lobulated cells had an increased propensity to arrest in G1. Indeed, G1 arrest is known to occur after paclitaxel treatment in non-transformed cell lines^31^.

To understand the relationship between transcriptomic variation, lobulation, and cell cycle, we examined the top batch-corrected principal components of the single cell transcriptomes. We noticed that the first four principal components separated cells by nuclear morphology (Supplementary Figure 4). To discover the genes associated with nuclear lobulation while controlling for the cell-free RNA batch effect, we sequenced an unseparated paclitaxel-treated cell population and aligned their transcriptomes to those from morphologically-normal and lobulated cells^32^. We then derived a lobulation score for each cell via linear combinations of the four principle components that correlate with nuclear morphology. Finally, we extracted genes associated with this lobulation score, which is higher in cells with lobulated nuclei, in the unseparated cells by using a differentially expressed gene test (Methods).

In total, 765 genes were significantly associated with the lobulation score (adjusted p-value < 0.01; Supplementary Figure 4, Supplementary Table 5). To our surprise, the vast majority (84%) of these genes were more highly expressed by morphologically normal cells. Morphologically normal cells upregulated the genes encoding actin and microtubules (e.g. *ACTB*, *TUBB4B*; Figure 4E), a well-documented response to microtubule damage and paclitaxel treatment^33^. We also noted that these cells upregulated the chaperone clusterin (*CLU*) and its co-activator *HSPA5*, which together decrease paclitaxel-mediated apoptosis by stabilizing mitochondrial membrane potential^34^. Intrigued by the notion that morphologically normal cells are resisting the effects of paclitaxel, we searched the literature for other genes upregulated in these cells and found that many of them, including *PRMT1*^35^*, ENO1*^36^*, STMN1*^37^*, LDHA*^38^*, ANXA5*^39^, and *HSPA8*^40^, are associated with paclitaxel resistance in diverse cancers.

To better understand the gene expression program associated with normal nuclear morphology in the context of paclitaxel treatment, we looked for enrichment of genes in previously defined gene sets^41^ covering a host of cellular processes (Supplementary Tables 6, 7). Morphologically normal cells upregulated 7 out of 8 proteins in the chaperonin containing TCP-1 complex (adjusted p value = 7.64e-15; Figure 4E), which is critical for tubulin folding and has been previously associated with paclitaxel resistance in ovarian cancer^39^. Morphologically normal cells also upregulated the transcriptional targets of two pacltiaxel resistance-associated signaling pathways^42, 43^: c-Myc (adjusted p value = 1.66e-30) and mTORC1 (adjusted p value = 6.19e-17; Figure 4F).Together, these results suggest that the morphologically normal, paclitaxel-treated cells mounted a biosynthetic and proteostatic response to drug treatment, with remarkable similarities to the gene expression profiles observed in paclitaxel resistant cell lines and cancers.

## Discussion

A major limitation of current microscopy-based experiments is the inability to isolate hundreds of thousands of phenotypically defined cells for further analysis. We developed Visual Cell Sorting, a microscope-based method that directs a digital micromirror device to irreversibly photoactivate a genetically encoded fluorescent protein in cells of interest, effectively translating a complex visual phenotype into one that can be sorted by FACS.

To highlight the Visual Cell Sorting’s flexibility, we performed two distinct experiments. First, we leveraged its high throughput to quantify the function of hundreds of nuclear localization sequence variants in a pooled, image-based genetic screen. By combining single variants that individually improved NLS function, we created an eight-residue superNLS (PPRKKRKI) that could be used to improve CRISPR-mediated genome editing, fluorescent protein-based nuclear labelling, and other experiments that leverage nuclear recombinant proteins. We then used the variant scores to make an accurate, amino acid preference-based predictor of NLS function in the human nuclear proteome. Second, we leveraged Visual Cell Sorting’s ability to recover live, phenotypically defined subsets of cells to investigate the heterogenous cellular response to paclitaxel treatment using single cell RNA sequencing. To our surprise, cells that resist the effect of paclitaxel on nuclear morphology appear to be adapting to its effects at the molecular level using a gene expression program with many similarities to paclitaxel-resistant cancers.

High throughput is a key advantage of Visual Cell Sorting, compared to other similar methods. In our pooled image-based screen, we analyzed approximately one million cultured human cells across 60 hours of imaging and sorting time, ultimately recovering ∼650,000. This throughput is ∼1,000-fold more than what could be achieved using other photoconvertible fluorophore-based methods^10–13^, ∼20-fold more than current MERFISH pooled screens^5^, and similar in per-day throughput to *in situ* sequencing-based screens^16^ (Supplementary Table 1). Thus, Visual Cell Sorting enables the analysis of thousands of genetic variants in a single experiment. Visual Cell Sorting throughput could be increased even further by analyzing cellular phenotypes at a lower magnification, by applying faster image analysis algorithms, or by shutting off Dendra2 expression before imaging to extend imaging time (Supplementary Figure 1).

A second key advantage of Visual Cell Sorting is that it does not require any expensive dye-based reagents such as oligo libraries or fluorescent-labelled oligos; customized hardware components; or complex workflows. Outfitting an automated wide-field microscope requires just three inexpensive, commercially available components: a live cell incubation chamber, a digital micromirror device, and a 405 nm laser. Finally, Visual Cell Sorting enables recovery of cells with up to four distinct phenotypes in one experiment, unlike other photoconvertible fluorophore-based methods^10–14^.

Visual Cell Sorting has important limitations. Cells must be genetically engineered to express Dendra2, which is photoactivated by blue fluorescent protein (BFP) excitation wavelengths and emits at GFP and RFP wavelengths. This requirement limits the other fluorescent channels are available for imaging. However, miRFP^44^ and mBeRFP^45^ can be used in conjunction with Dendra2, allowing two additional compartments or proteins to be marked in each experiment. Moreover, new analytical approaches leveraging brightfield images may reduce the need for fluorescent markers^46, 47^. Another limitation is that, unlike morphological profiling approaches^48^, Visual Cell Sorting requires a pre-defined phenotype of interest. Finally, Visual Cell Sorting experiments are limited to approximately twelve hours to avoid Dendra2 activation signal decay or cell overgrowth. However, the workflow we present, with imaging at 20X magnification and image processing times of 3-8 seconds, is sufficient for the analysis of hundreds of thousands of cells in each experiment.

In summary, Visual Cell Sorting is a robust and flexible method that can be used to separate heterogeneous cultures of cells into up to four morphologically defined subpopulations. The components required for Visual Cell Sorting are already in widespread use, are commercially available and can be adapted to most modern automated widefield fluorescent microscopes. The method will improve in scope and speed as further advances are made in cell segmentation and image analysis. We demonstrate that Visual Cell Sorting can be used for both image-based pooled genetic screens and image-based transcriptomics experiments. This flexibility should drive the application of Visual Cell Sorting to a wide range of biological problems in diverse fields of research that seek to dissect cellular heterogeneity, including stem cell biology, functional genomics, and cellular pharmacology.

## Methods

### General reagents, DNA oligonucleotides and plasmids

Unless otherwise noted, all chemicals were obtained from Sigma and all enzymes were obtained from New England Biolabs. KAPA Hifi 2x Polymerase (Kapa Biosystems KK2601) was used for all cloning and library production steps. *E. coli* were cultured at 37 °C in Luria broth. All cell culture reagents were purchased from ThermoFisher Scientific unless otherwise noted. HEK 293T cells (ATCC CRL-3216) and U-2 OS cells (ATCC HTB-96), and derivatives thereof were cultured in Dulbecco’s modified Eagle’s medium supplemented with 10% fetal bovine serum, 100 U/mL penicillin, 0.1 mg/mL streptomycin, and 1 ug/mL doxycycline (Sigma), unless otherwise noted. hTERT RPE-1 cells (ATCC CRL-4000) and derivatives thereof were cultured in F12/DMEM supplemented with 10% FBS, 1 mM PenStrep, and 0.01 mg/mL hygromycin B. For Visual Cell Sorting experiments, DMEM without phenol red was used to reduce background fluorescence. Cells were passaged by detachment with trypsin–EDTA 0.25%. All cell lines tested negative for mycoplasma. All synthetic oligonucleotides were obtained from IDT and their sequences can be found in Supplementary Table 8. All non-library-related plasmid modifications were performed with Gibson assembly. See the Supplementary Note for construction of the vectors used.

### Construction of the SV40 NLS Library

A site saturation mutagenesis library of the SV40 NLS upstream of a tetramerizing miRFP reporter (attB-NLS-CMPK-miRFP library) was constructed using Gibson cloning^49^. See the Supplementary Note for a detailed description of the construction of the site-saturation mutagenesis library.

### Cell Lines

U-2 OS cells expressing the Tet-ON Bxb1 landing pad (U-2 OS AAVS-LP Clone 11) were generated as previously described^50^. To create H3-Dendra2- and H3-Dendra2/H2B-miRFP-expressing derivative cell lines, attB-H3-Dendra2 or attB-H3-Dendra2-P2A-H2B-miRFP703 were recombined into U-2 OS AAVS-LP Clone 11 cells, as previously described^50^. For the NLS work, a separate clonal U-2 OS cell line expressing the Tet-ON landing pad and CMV-H3-Dendra2 was created by co-transduction of parental U-2 OS cells with the LLP-Blast lentivirus^51^ and another expressing histone H3-Dendra2 (U-2 OS LLP-Blast/H3-Dendra2 Clone 4). A clonal hTERT RPE-1 cell line expressing CMV-NLS_SV40_-Dendra2-GSSG-Dendra2-GSSG-Dendra2 (NLS-Dendra2×3); CMV-H2B-miRFP; and CMV-NES-mBeRFP was generated by transduction of a parental line (ATCC, CRL-4000) with three lentiviral vectors followed by single cell sorting (RPE-1 NLS-Dendra2×3/H2B-miRFP/NES-mBeRFP Clone 3). For more information regarding these lines and for the lentiviral production protocol, see the Supplementary Note.

### Recombination of single-variant SV40 NLS clones or the library into U-2 OS LLP-Blast/H3-Dendra2 Clone 4 cells

The SV40 NLS variant library or single-variant clones were recombined into U-2 OS LLP-Blast/H3-Dendra2 Clone 4 cells, as previously described in HEK 293Ts^50^. Two recombination replicates were performed. For more information, see the Supplementary Note.

### Visual Cell Sorting: equipment and settings

A Lecia DMi8 Inverted Microscope was outfitted with Adaptive Focus; an Incubator i8 chamber with PeCon TempController 2000-1 and Oko CO2 regulator set to 5%; a 6-line Lumencor Spectra X Light Engine LED; Semrock multi-band dichroic filters (Spectra Services, cat. no. LED-DA-FI-TR-Cy5-4X-A-000, LED-CFP/YFP/mCherry-3X-A-000); BrightLine bandpass emissions filters for DAPI (433/24 nm), GFP (520/35 nm), RFP (600/37 nm), NIR (680/22 nm), CFP (482/25 nm), YFP (444/24 nm), and mCherry (641/74 nm); a 20X 0.8 NA apochromatic objective; and a Mosaic3 Digital Micromirror Device affixed to a Mosaic SS 405 nm/1.1 W laser and mapped to an Ixon 888 Ultra EMCCD monochrome camera. The microscope and digital micromirror device were controlled with the Metamorph Advanced Image Acquisition software package (v7.10.1.161; Molecular Devices). The image size was ∼560 × 495 μm. Image bit depth ranged from 12-16 bits, depending on the brightness of cells in the field of view.

### Visual Cell Sorting: imaging, analysis and photoactivation

Cells were imaged on glass-bottom, black-walled plates (CellVis P06-1.5H-N, P24-1.5H-N, P96-1.5H-N) in phenol-red free media at 5% CO_2_ and 37 °C using the 20X 0.8 NA objective. ∼50-100 cells were imaged per field of view. To image unactivated Dendra2, 474/24 nm excitation and 482/25 nm emission filters were used. To image activated Dendra2, 554/23 nm excitation and 600/37 nm emission filters were used. To image miRFP, 635/18 nm excitation and 680/22 nm emission filters were used. Prior to imaging, the Auto Focus Control system was activated. Metamorph’s Plate Acquisition module was used to collect images and run Metamorph jounals that analyzed cells and directed their selective photoactivation by the digital micromirror device. For more information about the Metamorph journals used to image and activate cells, see the Supplementary Note.

### Visual Cell Sorting: FACS on Microscope-Activated Cells

Cells activated on the microscope were analyzed using an LSR II (BD Biosciences) or sorted into bins according to their Dendra2 photoactivation state using a FACS Aria III (BD Biosciences). First, live, single, Dendra2-positive cells were selected first using forward and side scatter, and then the FITC channel. Cells with different levels of photoactivation (e.g. no photoactivation, 50ms, 200ms, and 800ms) were gated using a PE-YG (activated Dendra2) vs. FITC (unactivated Dendra2) two-dimensional plot. Then, a PE-YG:FITC ratiometric parameter in the BD FACSDIVA software was created. A histogram of the FITC:PE-YG ratio was created and gates dividing cells into up to four bins was performed by using the populations identified on the PE-YG vs. FITC plot. Data was analyzed using FlowCytometryTools (v0.5.0) in Python (v3.6.5) or flowCore (v1.11.20) in R (v3.6.0); raw .fcs files and code are available on GitHub.

### Selective photoactivation of cells expressing miRFP

U-2 OS AAVS-LP Clone 11 cells with attB-H3-Dendra2 or attB-H3-Dendra2-P2A-H2B-miRFP recombined into the landing pad were counted and mixed in ratios ranging from 0.5% to 50% miRFP-expressing cells, then 40,000 cells of each mixture were seeded into three wells of a 24-well plate. The next day, cells were placed on the microscope and imaged, analyzed, and activated at 661 sites across each well of the plate, covering ∼95% of the total well area. At each site, Dendra2 and miRFP were imaged with 2×2 binning; Metamorph’s Count Nuclei module was used on the miRFP image to identify miRFP-expressing cells; and a binary with regions corresponding to miRFP-expressing cells was passed to the digital micromirror device, which subsequently activated the cells. Once all sites were imaged, analyzed, and activated, the cells were subject to flow cytometry to assess unactivated Dendra2, activated Dendra2, and miRFP expression. The experiment was repeated two additional times for a total of three replicates. For the Metamorph journals used to analyze and activate cells, see the GitHub repository. For more information about the gating scheme used for this experiment, see the Supplementary Note.

### Photoactivation of cells for 0, 50, 200, and 800 milliseconds

U-2 OS AAVS-LP Clone 11 cells with attB-H3-Dendra2-P2A-H2B-miRFP recombined into the landing pad were seeded at 50,000 cells per well in a 6-well glass bottomed plate. The next day, cells were imaged for unactivated Dendra2 and miRFP at 100 sites (10×10 square) and quartiles of total miRFP intensity were measured using Metamorph. Then, cells across 661 sites in two wells were left unactivated or activated for 50ms, 200ms, or 800ms according to the miRFP intensity quartile to which they belonged (Q1 = 0-3803, Q2 = 3804-5839, Q3 = 7396-9674, Q4 = 9674+). For the Metamorph journals used to analyze and activate cells, see the GitHub repository.

### Testing for photoactivation-induced toxicity with Annexin V and DAPI

U-2 OS AAVS-LP Clone 11 cells with attB-H3-Dendra2 recombined into the landing pad were seeded at 20,000 cells per well in a 24-well plate. Over the next two days, cells across 400 sites (60% well coverage) in three replicate wells were segmented using the Count Nuclei module in Metamorph and activated for 800 ms. Forty-eight hours after the first well was activated, cells were trypsinized, stained with Annexin V (Thermo, cat. no. A23204) and DAPI (Invitrogen, cat. no. D1306), and subjected to flow cytometry to assess unactivated Dendra2, activated Dendra2, Annexin V, and DAPI. Three wells of unactivated cells were heated at 50 °C for 10 minutes as a cell death positive control. The experiment was repeated two additional times for a total of three replicates. Data was analyzed using FlowJo (v10.5.3).

### Testing for photoactivation-induced toxicity with RNA sequencing

U-2 OS AAVS-LP Clone 11 cells with attB-H3-Dendra2 recombined into the landing pad were seeded at 20,000 cells per well in 8 wells of a 24-well plate. Eighteen hours later, cells across 6 wells (678 sites per well; ∼100% well coverage) were activated and then incubated for 0.5, 1.5, 2.5, 3.5, 4.5, or 6 hours (1 well each). Two wells were left unactivated. Dendra2 photoactivation was verified by flow cytometry, with the two unactivated samples were used as negative controls. Bulk RNA sequencing libraries were prepared as described previously (Cao et. al. 2017). Briefly, RNA was extracted from each sample using a Trizol/RNeasy Mini Kit (ThermoFisher 15596026, Qiagen 74104) then subjected to SuperScript IV First-Strand Synthesis (Thermo Fisher 18091050) and NEBNext Ultra II Directional RNA Second Strand Synthesis (NEB E7550), according to the manufacturer’s instructions. cDNA was then tagmented with Nextera Tn5 (Illumina FC-131-1024) and amplified/indexed by PCR with the NEBNext DNA Library Prep Kit (NEB E6040). Samples were sequenced using a NextSeq 500/550 75 cycle kit (Illumina TG-160-2005). Differential gene expression analysis of RNA sequencing data followed the standard DESeq2 workflow^52^. Briefly, differential gene expression testing was performed using a binary coding of photoactivation status in the DESeq2 design formula. Dispersion estimates, log2 fold changes and adjusted p-values were all calculated using the DESeq() function with default parameters as specified in DESeq2.

### Visual Cell Sorting of cells expressing SV40 NLS library

Eighteen hours before imaging, 300,000 U-2 OS LLP-Blast/H3-Dendra2 Clone 4 cells with the attB-NLS-CMPK-miRFP library recombined into the landing pad were seeded into each well of a 6-well plate. The next day, cells were placed onto the microscope and imaged, analyzed, and activated across 2,949 sites (∼100% well coverage) across two wells. At each site, Dendra2 and miRFP were imaged with 2×2 binning; Metamorph’s Count Nuclei module was used on the Dendra2 image to identify nuclei and create a nuclear binary image; cytoplasm binaries were created by subjecting the nuclear binary to a dilate function and subtracting away the nuclear binary; each nucleus-cytoplasm binary pair was superimposed on the miRFP image and average pixel intensities were measured for each compartment; cells with an average nuclear or cytoplasmic miRFP pixel intensity of less than 11,000 were filtered out; a nucleus-to-cytoplasm (N:C) ratio was calculated by dividing the average nuclear pixel intensity by the average cytoplasmic pixel intensity; nuclei with N:C < 0.964 were not activated at all, N:C 0.964 – 1.079 were activated for 50 ms, N:C 1.079 – 1.244 were activated for 200 ms, and N:C > 1.244 were activated for 800 ms. Once all sites were imaged, analyzed, and activated, the cells were subject to FACS and unactivated Dendra2 (FITC), activated Dendra2 (PE-YG), and miRFP (AlexaFluor-700) fluorescence intensities assessed. Cells were then sorted into four photoactivation bins (Supplementary Figure 2). A total of two Visual Cell Sorting replicates were performed on recombination replicate 1, and three were performed on recombination replicate 2. The details of replicate sorts for the NLS library can be found in Supplementary Table 2.

### Sorted SV40 NLS library genomic DNA preparation and sequencing

After sorting, cells in each Dendra2 photoactivation bin were grown in the absence of doxycycline until confluent in one well of a 6-well plate (∼ 7 days), then pelleted and stored at −20 °C. DNA was extracted from cell pellets with the DNEasy kit (Qiagen 69504) using RNAse according to the manufacturer’s instructions. gDNA was amplified using SV40_NLS_seq_f and SV40_NLS_seq_r (Supplementary Table 8) primers using Kapa Hifi (Kapa Biosystems KK2602) according to the manufacturer’s instructions. Amplicons were cleaned using Ampure XP beads (Beckman Coulter A63880), then subjected to an indexing PCR step using KAPA2G Robust (Kapa Biosystems KK5705) with primers P5 and an indexing primer (Supplementary Table 8). Amplicons were then run on a 1.5% agarose gel at 130 V for 40 min and the DNA in the 235bp band extracted using Freeze’N Squeeze DNA Gel Extraction Spin Columns (BioRad 7326165). Extracted DNA was sequenced on an Illumina NextSeq500 using SV40_NLS_Read1, SV40_NLS_Read2, and SV40_NLS_Index1 primers (Supplementary Table 8). Reads were trimmed and merged using PEAR^53^. Sequences were quality-filtered and variants were called and counted by using Enrich2, as previously described^21^. The Enrich2 configuration file is available on the GitHub repository.

### Calculating NLS variant localization scores

Jupyter v5.5.0 running Python v3.6.5 was used for analyses of the Enrich2 output. First, two filters were applied to remove low-quality variants : (1) a minimum nucleotide variant count cutoff of 5 in each bin in each replicate and (2) a requirement that the variant was accessible via NNK codon mutagenesis. After filtering, remaining nucleotide variants encoding the same amino acid substitution were added to yield a sum of counts for that variant within each bin for each replicate. To generate raw quantitative scores (*S_raw_*), a weighted average approach as previously described^54^ was applied to the variant frequencies (*f_var_*) across the 4 bins (*b1* – *b4*) in each replicate:

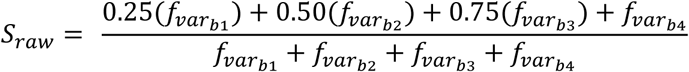

Raw scores were subsequently normalized such that variants with a wild-type raw score (*S_WT_*) have a normalized score of 1 and variants with the median raw score of the bottom 10% of variants (*S_P10_*) have a normalized score of 0:

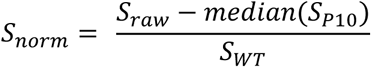

A final round of frequency filtering for variants, which sought to increase score correlations without excluding too many variants, removed variants present at a frequency lower than 0.003% of reads in all bins. Then, the raw and normalized scores were recalculated for each replicate; and the mean and standard error of the normalized scores from the five replicates were calculated to produce final scores. An iPython notebook file with the code used to run the analysis is available on the GitHub repository.

### Validation of single NLS variants

ssDNA oligos (IDT) encoding the NLS variants were introduced into EcoRI-digested attB-EcoRI-CMPK-miRFP reporter plasmid via a Gibson reaction^49^. Variants were validated by Sanger sequencing. Plasmids were recombined into 80,000 U-2 OS cells in a 24 well plate using 1.5 uL of FuGENE6 (Promega E2691) in 100 uL OPTIMEM (Fisher Scientific 31985070) with 100 ng of pCAG-Bxb1 and 295 ng of the attB variant recombination plasmid. After 5 days, recombined cells, which are miRFP+, were isolated using FACS for miRFP+ cells and plated in glass-bottom 24 well plates. Then, recombined cells were imaged for H3-Dendra2 and miRFP. Metamorph was used to segment nuclei and calculate mean nuclear and cytoplasmic miRFP intensity for each cell, as described above (“Visual Cell Sorting on cells expressing SV40 NLS library”). miRFP intensities were background-corrected (see Supplementary Note), and cells with nuclear and cytoplasm miRFP intensities roughly equal to background levels were removed. Then, N:C ratios were calculated for each cell using the cell’s mean nuclear (*I_nu_*_c_) and cytoplasmic (*I_cyt_*) miRFP intensities:

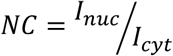

Each variant was examined in at least three separate imaging replicates. For more information regarding the validation of single NLS variants, see the Supplementary Note.

### Prediction of novel human NLS’s

Analysis of the normalized variant localization scores were done in RStudio v1.1.456 running R v3.6.0. Position-wise amino acid preferences were calculated^22^f:

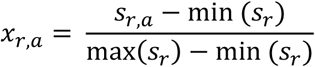

Where *x_r,a_* is the amino acid preference for amino acid *a* at position *r*, *s_r,a_* is the mean raw score of variants with amino acid *a* at position *r*, and *sr* is the set of all raw scores at position *r*. The scores of missing variants were estimated using the median score at that variant’s position. To train a weighted preference model, NLS sequences (n = 573) were downloaded from UniProt using a SPARQL query for all human proteins with a sequence motif annotation that contained the string “Nuclear localization” in its comment. A set of 573 “likely NLS” 11mers were generated by repeating the following for each NLS: (1) scoring every 11mer peptide overlapping the annotated NLS sequence by summing the amino acid preferences of the 11mer peptide (2) annotating the maximum-scoring 11mer as a “likely NLS”. All other possible 11mers in the training dataset (333,255 total) were annotated as “no NLS”. To account for the fact that some the amino acid preferences at some positions may be more important than others, a linear regression model of the following form was fit to these data:

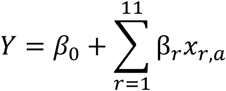

Where *Y* denotes the sequence class (“no NLS” = 0, “likely NLS” = 1), *β_0_* is the intercept, *β_r_* is the weight given to the amino acid preferences at position *r*, and *x_r,a_* is the is the preference of amino acid *a* at position *r*. Model parameters were determined by 8-fold cross validation before being applied to an independent test dataset^18^ containing 20 protein sequences with 30 NLS’s that were not examined during training.

To apply the final model to the nuclear human proteome, the test dataset was used to generate two score cutoffs: one corresponding to a precision of 0.9 (“high confidence NLS”) and one corresponding to a recall of 0.9 (“candidate NLS”). All 11mers present in proteins annotated as nuclear by the Human Protein Atlas were then subject to scoring by the model. An R-markdown file with the code used to run the analysis is available on the GitHub repository.

### Time-lapse imaging of cells treated with paclitaxel

hTERT-RPE-1 cells expressing Dendra2-NLS, H2B-miRFP703, and mBeRFP-NES were plated at a density of 50,000 cells per well in 2-well µm-slide chambers (iBidi). Twenty-four hours after plating, the cell media was replaced with media containing 0.25 nM taxol. After the cell media change, the cells were imaged for 24 hours with a pass time of 10 minutes. Imaging was performed on a Leica DMi8/Yokagawa spinning disk confocal microscope with a 20x 0.8NA air objective at 37°C and 5% CO2. Images were captured with an Andor iXon Ultra camera using Metamorph software. Videos were cropped and adjusted for brightness and contrast using ImageJ and Photoshop.

### Visual Cell Sorting of cells treated with paclitaxel

RPE-1 NLS-Dendra2×3/H2B-miRFP/NES-mBeRFP Clone 3 cells were plated at 50,000 cells per well in a 6-well plate. After 24 hours, cells were treated with paclitaxel at a final concentration of 0.25 nM. After 30 hours of treatment, cells were placed on the microscope and imaged, analyzed, and activated across 2,204 sites (∼75% coverage, avoided well edges) in 2 wells. At each site, Dendra2 was imaged with 1×1 binning; a custom nuclear segmentation pipeline that optimized detection of nuclear blebs, herniations, and other abnormalities was employed (see Supplementary Note); Metamorph’s MDA analysis was used to compute shape factors for nuclear binaries. Cells with nuclear shape factor < 0.65 were activated for 200 ms, and cells with nuclear shape factor > 0.65 were activated for 800 ms. Cells from each well were trypsinized and resuspended in DPBS supplemented with 1% BSA and 2% FBS. Using FACS, cells corresponding to 200 ms and 800 ms photoactivation were sorted using FACS (Supplementary Figure 3) into a 1.5 mL tube containing 1mL DPBS supplemented with 1% BSA. In Experiment 1, cells were sorted according to their nuclear phenotype (i.e. 200 ms cells in bin 1, 800 ms cells in bin 2; Fig S4A). Cells were imaged, activated, and sorted identically in Experiment 2, except that all activated cells were sorted into one bin (i.e. both 200 ms and 800 ms cells in bin 1; “unseparated paclitaxel-treated population”).

### Single cell RNA sequencing of sorted, paclitaxel-treated populations

After sorting, cells were spun at 1,000 × g at 4 °C for 5 minutes, then all but 50-100 uL of supernatant was removed. Cells were counted and subjected to 10X Single-Cell RNA sequencing v2 (10x Genomics, cat. no. 120236, 12037) according to the manufacturer’s instructions. 10x Cell Ranger version 2.1.1 was used to process lanes corresponding to the single cell libraries and map reads to the human reference genome build Hg19. Unique molecular identifier (UMI) cutoffs were chosen by 10x Cell Ranger software. Reads and cell numbers were normalized via downsampling by the aggregate function in 10x Cell Ranger. After normalization, cells had a median of 9,249 UMIs (Experiment 1, separated populations) or 16,932 (Experiment 2, unseparated population) per cell.

### Analysis of single cell RNA sequencing data

Analysis of 10X CellRanger output files was done in RStudio v1.1.456 running R 3.6.0. Cell cycle scoring and annotations were performed with Seurat, as previously described^30^. UMAP was performed with Monocle3^55, 56^. Mutual-nearest neighbors batch correction was performed using the Batchelor package^32^ in the following order: unseparated cells from Experiment 2 were batch corrected with morphologically-normal cells from Experiment 1, and then lobulated cells from Experiment 1 were batch corrected. An R-markdown file with the code used to run the analysis is available on the GitHub repository.

### Differentially Expressed Genes Analysis

Mutual nearest neighbors batch correction^32^ was used to align cells from Experiment 2 (normal and lobulated cells sorted into the same tube, one 10X lane) to cells from Experiment 1 (normal and lobulated cells sorted into separate tubes, two 10X lanes). Principle components 1 through 4, which were output by the batch correction algorithm, were used to train a logistic regression model for nuclear lobulation on the cells in Experiment 1. This model was applied to Experiment 2, resulting in each cell being assigned a lobulation score, which is high in lobulated cells in Experiment 1 and low in normal cells in Experiment 1. Then, a differentially expressed gene test was performed on the cells in Experiment 2 using lobulation score, Seurat-computed G1 score, and Seurat-computed G2/M score as covariates. For a detailed discussion of this analysis, see the Supplementary Note.

### Gene Set Enrichment Analysis

Gene set enrichment analysis was performed using the piano package^57^ in R on differentially expressed genes with a log2-normalized effect value (equivalent to the expected log2-fold change per unit increase in lobulation score) less than −0.1 and a q-value less than 0.01. The MSigDB Hallmarks and Canonical Pathways gene sets were used^41, 58^.

### Data availability

Illumina sequencing bam files, fastq files, final UMI matrices, the gene and cell annotations from Cell Ranger, and other large preprocessed data files can be accessed at the NCBI Gene Expression Omnibus (GEO) data repository under accession number GSE141030. The data repository will be made publicly available at the time of publication.

### Code availability

Preprocessed data and code used to generate the figures and tables in this manuscript is available with permission at the Fowler lab GitHub repository (https://github.com/FowlerLab/vcs_2019.git). The code will be made publicly available at the time of publication.

## Acknowledgements

Kate Sitko, Gabriel Boyle, and Kathleen Abadie contributed to the development of Visual Cell Sorting and helped refine its capabilities. Jose McFaline provided advice for the interpretation and analysis of single cell RNA sequencing results. Sri Kosuri provided valuable insight and encouragement. Stanley Fields and Brian Beliveau provided helpful comments on the manuscript. This work was funded by the NIH (F30CA236335-01 to N.H., R01GM109110 and RM1HG010461 to D.M.F., R35GM124766-03 and P30CA015704 to E.M.H, DP2HD088158 to C.T., R01HL120948 to R.J.M Jr.), the W. M. Keck Foundation (C.T.), the NSF (DGE-1258485 to S.S), and The Paul G. Allen Frontiers Group (C.T.).

## Contributions

N.H. and D.M.F. conceived of Visual Cell Sorting. N.H. and A.C. carried out Visual Cell Sorting experiments. S.S., J.S. and D.J. prepared next-generation sequencing libraries. Z.K. validated NLS variant scores. S.P. cloned constructs and helped with preliminary experiments. E.M.H. guided the paclitaxel experiment. W.T. and R.J.M. Jr. created the U-2 OS AAVS-LP Clone 11 cells. E.M.H. and H.H. provided hTERT-RPE-1 cells and performed time-lapse imaging. S.S. and C.T. provided valuable insight for the analysis of single cell RNA sequencing results. N.H. and D.M.F. wrote the manuscript. All authors commented on the manuscript.

## Supplementary Figures

**Supplementary Figure 1.**
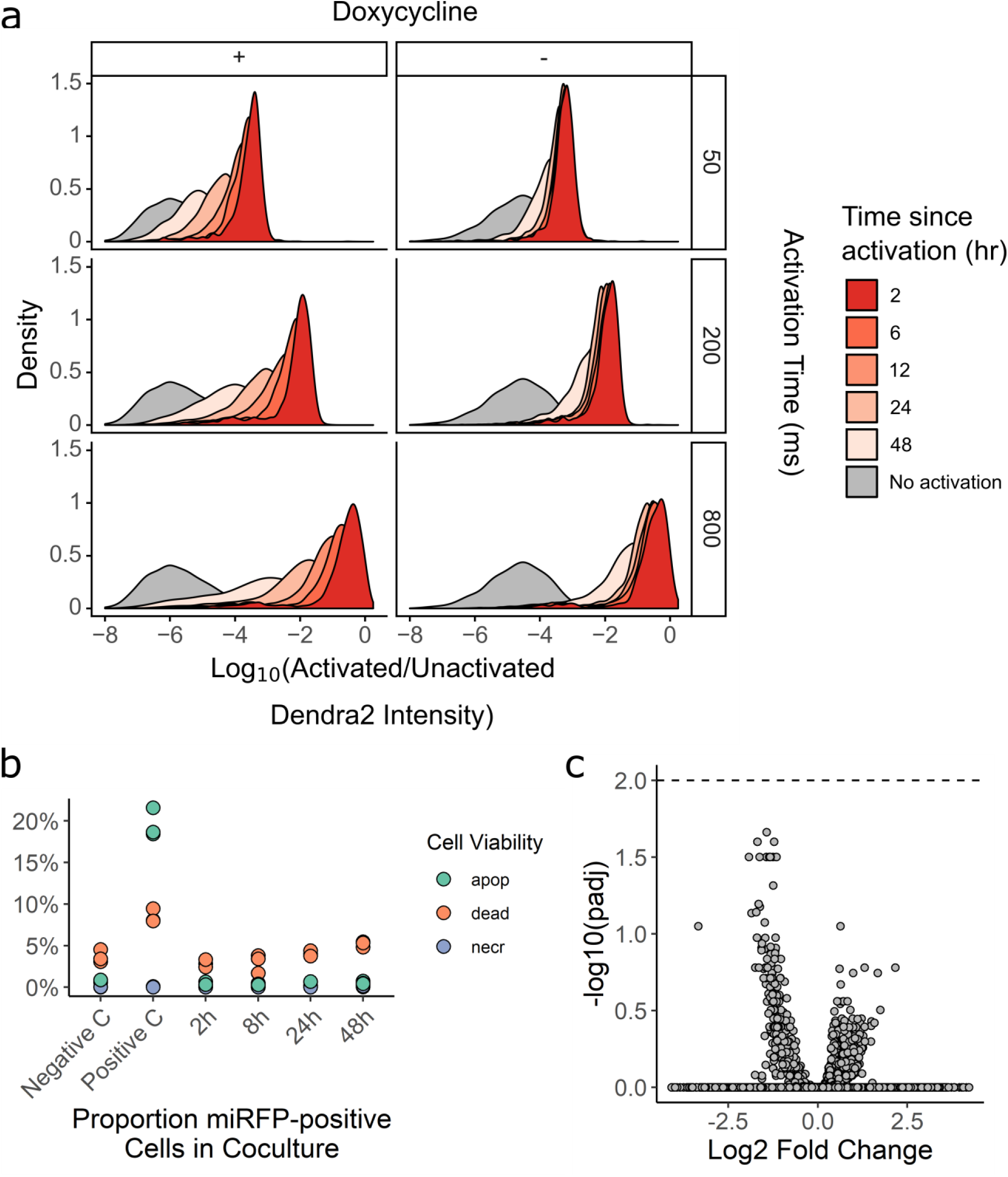
Visual Cell Sorting. **(a)** U-2 OS cells expressing H3-Dendra2 under the control of a doxycycline inducible promoter were activated with 405 nm light for 50, 200, or 800 ms; incubated for various lengths of time; and then subject to flow cytometry to determine the degree of activated Dendra2 (left panel). To examine whether shutting off Dendra2 expression before the experiment increases photoactivation ratio stability, the experiment was repeated, but doxycycline was removed from the media before cells were placed under the microscope (right panel) **(b)** To examine the effect of Dendra2 photoactivation on cell viability, cells were activated for 800 ms and then apoptosis, necrosis, and death were assessed by flow cytometry using DAPI and Annexin-V. Negative C, no photoactivation. Positive C, incubation of cells at 50 C for 10 min; apop, apoptotic cells; necr, necrotic cells (n. **(c)** To test whether Dendra2 photoactivation affects gene expression, cells were activated for 800ms, incubated for 0.5, 1.5, 2.5, 3.5, 4.5, or 6 hours and subsequently subject to bulk RNA seq. Samples were compared to two separate replicates of unactivated cells. Volcano plot of differentially expressed genes shown. Dotted line, adjusted p-value of 0.01.

**Supplementary Figure 2.**
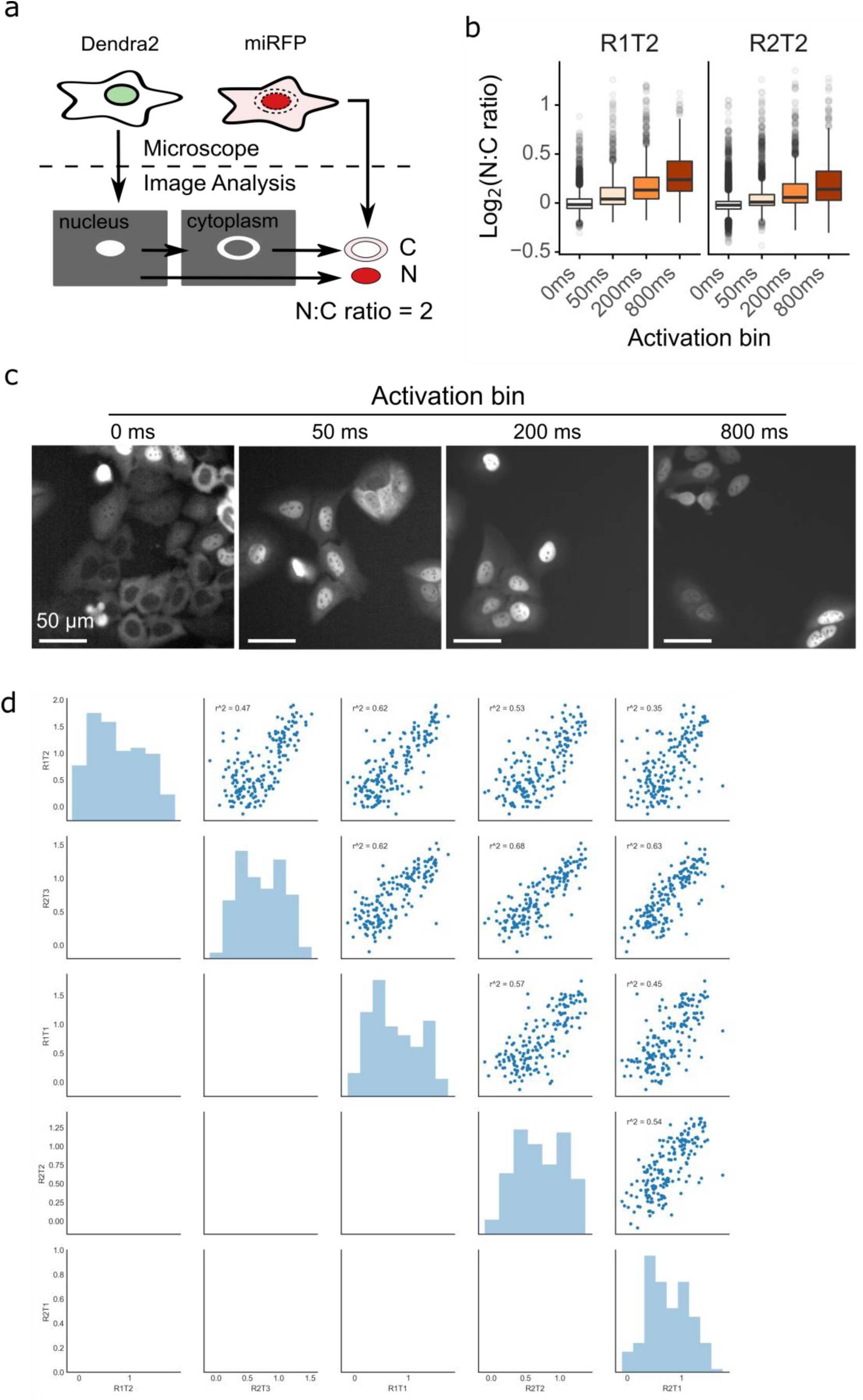
Visual Cell Sorting for pooled, image-based genetic screening. **(a)** Image analysis pipeline to calculate nucleus to cytoplasm (NC) ratio. Nuclei were segmented using the H3-Dendra2 signal. Cytoplasmic masks were created by dilating and then removing the nuclear mask. Mean miRFP intensity was measured within each mask and the nucleus to cytoplasm (N:C) ratio calculated. **(b)** After selective photoactivation on the microscope based on N:C ratio, cells were subject to fluorescence-activated cell sorting and sorted according to their nuclear localization phenotype. Two days after sorting, cells from each sort bin were re-imaged and the nucleus to cytoplasm ratio reassessed (n = ∼1,500 per photoactivation bin). Center line, median; box limits, upper and lower quartiles; whiskers, 1.5x interquartile range; points, outliers. **(c)** Representative images of sorted cells. **(d)** Correlation plots of normalized scores calculated for each replicate. r^2, square of Pearson’s correlation coefficient.

**Supplementary Figure 3.**
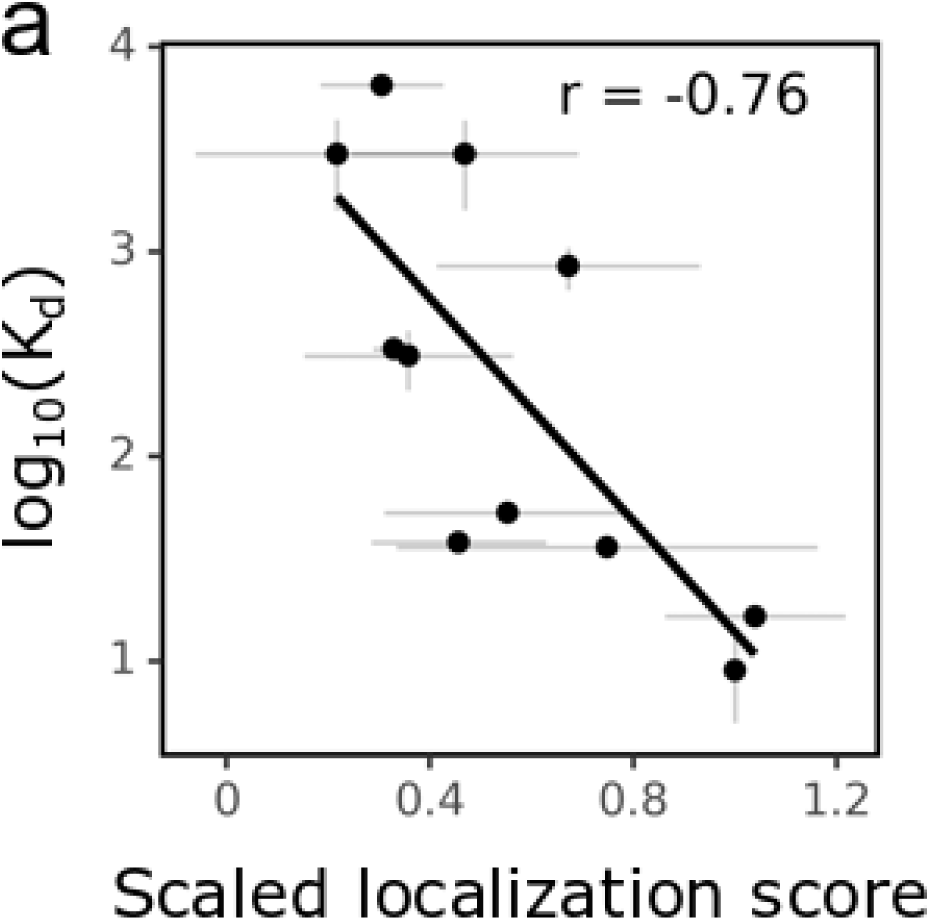
Visual Cell Sorting-derived variant scores accurately predict NLS function. **(a)** To examine whether localization score is related to binding to importin alpha, dissociation constants measuring binding between SV40 NLS variants and importin alpha, as reported by Hodel and colleagues (2001) were plotted against the variants’ mean normalized scores. Grey bars, standard error from the mean. r, Pearson’s correlation coefficient.

**Supplementary Figure 4.**
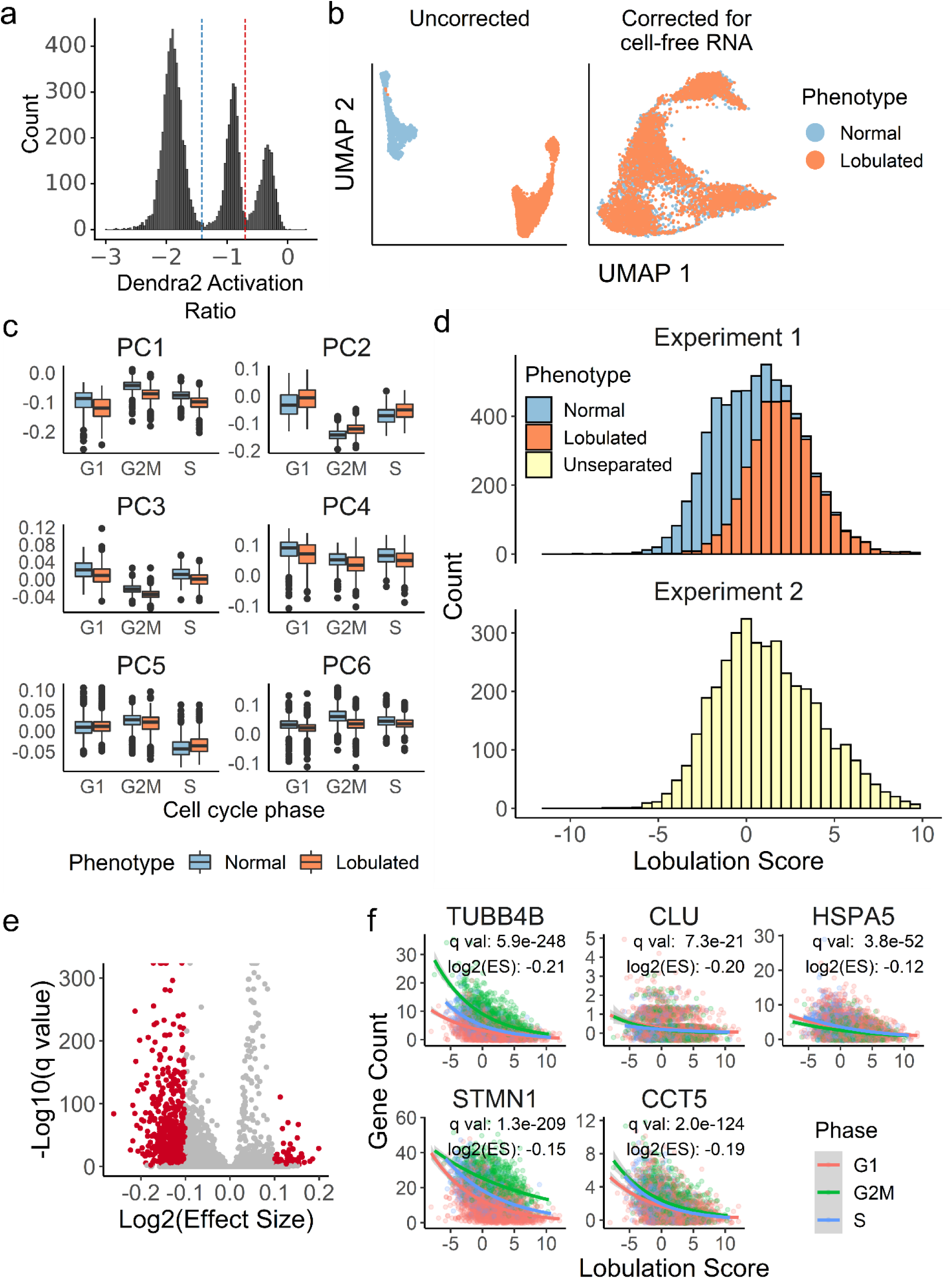
**(a)** Photoactivation gates for Visual Cell Sorting. Cells between the blue and red dotted lines represent putative normal nuclear shape factor cells activated with 405 nm light for 200 ms; and cells above the red dotted line represent putative low nuclear shape factor (lobulated) cells activated for 800 ms. Cells below the blue dotted line were not imaged or not activated **(b)** UMAP projection of the single cell transcriptomes derived from Visual Cell Sorting-separated lobulated and normal cells before and cell-free RNA correction with the algorithm used by SoupX^27^ **(c)** Visual Cell Sorting-separated cells were aligned to an unseparated, paclitaxel-treated population with the mutual nearest neighbors algorithm^32^. The first six principle components of the separated cells, subset by nuclear phenotype and cell cycle stage, are shown. **(d)** Lobulation scores were generated using linear combinations of principle components 1-4. Top, Visual Cell Sorting experiment (N = 6,277 cells); bottom, unseparated cell population (N = 3,859 cells. **(e)** Volcano plot showing DEGs significantly correlated with the lobulation score in the unseparated, paclitaxel-treated population. Red points, significant DEGs with a log2(Effect Size), which estimates the expected log2 fold-change per unit increase in lobulation score, greater than 0.1 and a q-value less than 0.01. **(f)** Raw gene counts of selected significant DEGs of cells in the unseparated, paclitaxel-treated population versus the cells’ lobulation scores. Colored lines, negative binomial regression model stratified by cell cycle stage; ES, effect size, the expected fold change in gene expression per unit increase in lobulation score.

## Supplementary Tables

**Supplementary Table 1.** Comparison of microscope-based methods that dissect the relationship between genetic perturbations or gene expression programs and visual phenotype.

**Supplementary Table 2.** Number of cells sorted for each replicate of the image-based screen of SV40 NLS variants.

**Supplementary Table 3.** SV40-NLS missense variant scores. Raw scores are equivalent to unscaled weighted averages of variant frequencies across sort bins. Scaled scores are normalized such that the intra-replicate wild-type score equals 1 and the median of the bottom 10% of scores equals 0.

**Supplementary Table 4.** Monopartite NLS predictions for nuclear human proteins, determined using a model that leverages the scaled scores of single SV40-NLS variants.

**Supplementary Table 5.** Differentially-expressed genes that are significantly associated with the lobulation score metric in unseparated, paclitaxel-treated cells. Lobulation scores were derived by batch-correcting unseparated, paclitaxel-treated cells with paclitaxel-treated cells sorted according to their degree of nuclear lobulation.

**Supplementary Table 6.** MSigDB Cellular Process gene sets that are significantly enriched in cells with low lobulation scores (i.e. morphologically normal cells).

**Supplementary Table 7.** MSigDB Hallmarks gene sets that are significantly enriched in cells with low lobulation scores (i.e. morphologically normal cells).

**Supplementary Table 8.** Primers and plasmids used for this manuscript.

## Supplementary Videos

**Supplementary Videos 1 and 2.** Live cell imaging of Dendra2 at 20X magnification in asynchronous hTERT RPE-1 expressing NLS-Dendra2×3 H2B-miRFP70.3 cells treated with

0.25 nM paclitaxel for 24 hours. Each second in the videos is 1.2 hours.

## Supplementary Note

### Extended description of general reagents, DNA oligonucleotides and plasmids

To create attB-H3-Dendra2, the Dendra2 open reading frame was obtained from Dendra2-Lifeact7 (a gift from Michael Davidson Addgene #54694) and cloned downstream of the H3 open reading frame from mEmerald-H3-23 (a gift from Michael Davidson Addgene #54115) and into the backbone of attB-EGFP-PTEN-IRES-mCherry^50^.

To create attB-H3-Dendra2-P2A-H2B-miRFP703, attB-H3-Dendra2 and pH2B-miRFP703 (a gift from Vladislav Verkhusha, Addgene #80001) were combined and a P2A sequence included in the Gibson overhang regions between Dendra2 and miRFP.

To create pLenti-CMV-H3-Dendra2, the H3-Dendra2 reading frame in attB-H3-Dendra2 replaced the open reading frame in pLenti CMV rtTA3 Blast (w756-1), a gift from Eric Campeau (Addgene #26429).

To create attB-Nterm-CMPK-miRFP (the destination vector for the NLS library), a gBlock encoding an EcoRI site in-frame and upstream of CMPK (IDT; based off a previously published SV40 NLS construct^20^) was combined with the miRFP open reading frame from pH2B-miRFP703 and inserted into the backbone of attB-H3-Dendra2-P2A-H2B-miRFP703.

To create attB-NLS-CMPK-miRFP and all single, double, and triple amino acid variants, the attB-Nterm-CMPK-miRFP vector was digested with EcoRI for 2 hours at 37 C. Then, the digested plasmid and an oligo that contained the NLS (wild-type or variant of interest) and 55 C overhangs complementary to the edges of the cut site were incubated in a Gibson reaction in a one to three molar ratio and transformed, as per manufacturer’s instructions.

To create pLenti-CMV-NLS-Dendra2×3-P2A-H2B-miRFP, three PCRs of Dendra2 (template derived from Dendra2-Lifeact7) were performed: one with an N-terminal NLS appended on the forward primer and a Gly-rich linker on the reverse; one with the Gly-rich linker on the forward primer and a second, non-identical Gly-rich linker on the reverse primer; and one with the second Gly-rich linker on the forward primer and a stop codon on the reverse primer. These were combined with an attB construct backbone^50^ to create attB-NLS-Dendra2×3. In a second cloning step, H2B-miRFP from pH2B-miRFP703 was appended downstream to create attB-NLS-Dendra2×3-P2A-miRFP. Finally, the Dendra2×3-P2A-H2B-miRFP open reading frame was cloned into pLenti CMV rtTA3 Blast (w756-1).

To create pLenti-CMV-mBeRFP-NLS, a gBlock encoding codon-optimized mBeRFP^45^ (IDT) was cloned into pLenti CMV rtTA3 Blast (w756-1) with an NES encoded into Gibson overhangs.

### Extended description of the construction of site-saturation mutagenesis library for the SV40 NLS

The library of all possible SV40 NLS missense variants was constructed using a Gibson cloning approach. Eleven primer pairs – 1 for each NLS codon, plus 2 codons upstream and 2 codons downstream of the NLS – were designed (Supplementary Table 8). For each pair, the forward primer contained a 3’ annealing region (Tm ∼55C), an NNK codon, and a 5’ Gibson homology region (Tm ∼55C). The reverse primer comprised of the reverse complement of the forward primer Gibson homology region. Each primer pair was used in a separate PCR reaction that included attB-NLS-CMPK-miRFP as the template, and 5ul of each reaction were run on a 1% gel to check for product. The remaining 20ul was DpnI digested for 2 hours at 37 C to remove template plasmid, cleaned using DNA Clean & Concentrator-5 (Zymo Research D4013), subject to a 1-piece Gibson reaction, and transformed into chemically competent *E. coli*. Bulk transformant cultures were grown overnight and harvested using GenElute HP Plasmid DNA Midiprep Kit (NA0200-1KT). DNA preps containing single codon variant were subsequently mixed such that each prep contributed an equal amount of DNA to the final library.

### Extended description of lentivirus production

To produce lentivirus, HEK293T cells were plated in clear plastic 6 well plates (VWR, cat. no. 10062-892) at 4.5e5 cells per well. The next day, cells in each well were transfected with 1,125 ng psPAX2 (a gift from Didier Trono, AddGene #12260), 375 ng pMD2.G (a gift from Didier Trono, AddGene #12259), and 1,500ng of pLenti transfer vector using 6ul of FuGENE6 (Promega, cat. no. E2691) according to manufacturer’s instructions. Media was replaced 24 hours after transfection and collected at 48 hours and 72 hours after transfection. Collected media was spun at 1000g for 5 minutes, then the viral supernatant was decanted and filtered using a 0.45um filter (VWR, cat. no. 28145-481). Finally, the virus was concentrated using PEG-it Virus Precipitation Solution (SBI, cat. no. LV810A-1) and stored at −80C.

### Extended description of the creation of clonal cell lines

To create the clonal U-2 OS landing pad and H3-Dendra2 expressing line, parental U-2 OS cells were transduced with lentivirus encoding the landing pad^50^and lentivirus encoding H3-Dendra2. Five days after transduction, BFP +ve / Dendra2 +ve cells were sorted using an Aria III (Pacific Blue and FITC Channels). Three days later, cells were sorted directly into 96 well plates containing 75ul of conditioned U-2 OS media. Every week, 50ul of normal media was added to the well. Wells were checked for surviving clones at 2 weeks and 3 weeks post-sorting.

To create the hTERT RPE-1 clonal line expressing NLS-Dendra2×3, mBeRFP-NES, and H2B-miRFP, lentiviruses encoding these constructs were added to the parental line, and single cell clones were similarly sorted and expanded in conditioned media in 96 well plates.

### Extended description of the recombination of single-variant SV40 NLS clones or the library into the U-2 OS-landing pad line expressing H3-Dendra2

To recombine NLS variants or the NLS library into cells, H3-Dendra2 expressing U-2 OS cells with the landing pad were subject to Lipofectamine 3000 (Thermo Fisher L3000015) transfections in 6 well plates, T-25 flasks, or T-75 flasks, according to manufacturer instructions, with the following specifications: plated cells at 0.1e5 cells/well (24 well plate), 0.6e5 cells/well (6 well plate), 1.4e6 cells/flask (T-25), or 4.2e6/flask (T-75); transfected with 0.75ul/3.75ul/10.4ul/31.2ul Lipofectamine 3000, 1ul/5ul/13.9ul/41.7ul P3000 reagent, 500ng/2500ng/7000ng/21000ng total DNA at a by-weight ratio of 1/3 pCAG Bxb1 and 2/3 attB plasmid(s). Cells were transfected immediately after plating. Twenty-four hours after transfection, media was replaced. Doxycycline was added 48h after transfection. BFP negative, miRFP positive, Dendra2 positive cells were sorted 5-8 days after transfection.

### Extended description of the Metamorph journals used for imaging, analysis, and photoactivation

Visual Cell Sorting experiments have three Metamorph journals specified in the Metamorph high-throughput acquisition dialog box: a startup journal that initializes global variables accessed by other journals; an after-image journal that analyzes and activates cells; and an end of plate journal that turns off the laser. The microscope was directed to leave no overlap between images. In all experiments, nuclei touching the image border were removed. Site maps were customized by altering the htacquir.cfg configuration file. See the GitHub repository for the Metamorph journals and configuration files used.

### Extended description of the gating scheme for selective photoactivation of cells expressing miRFP

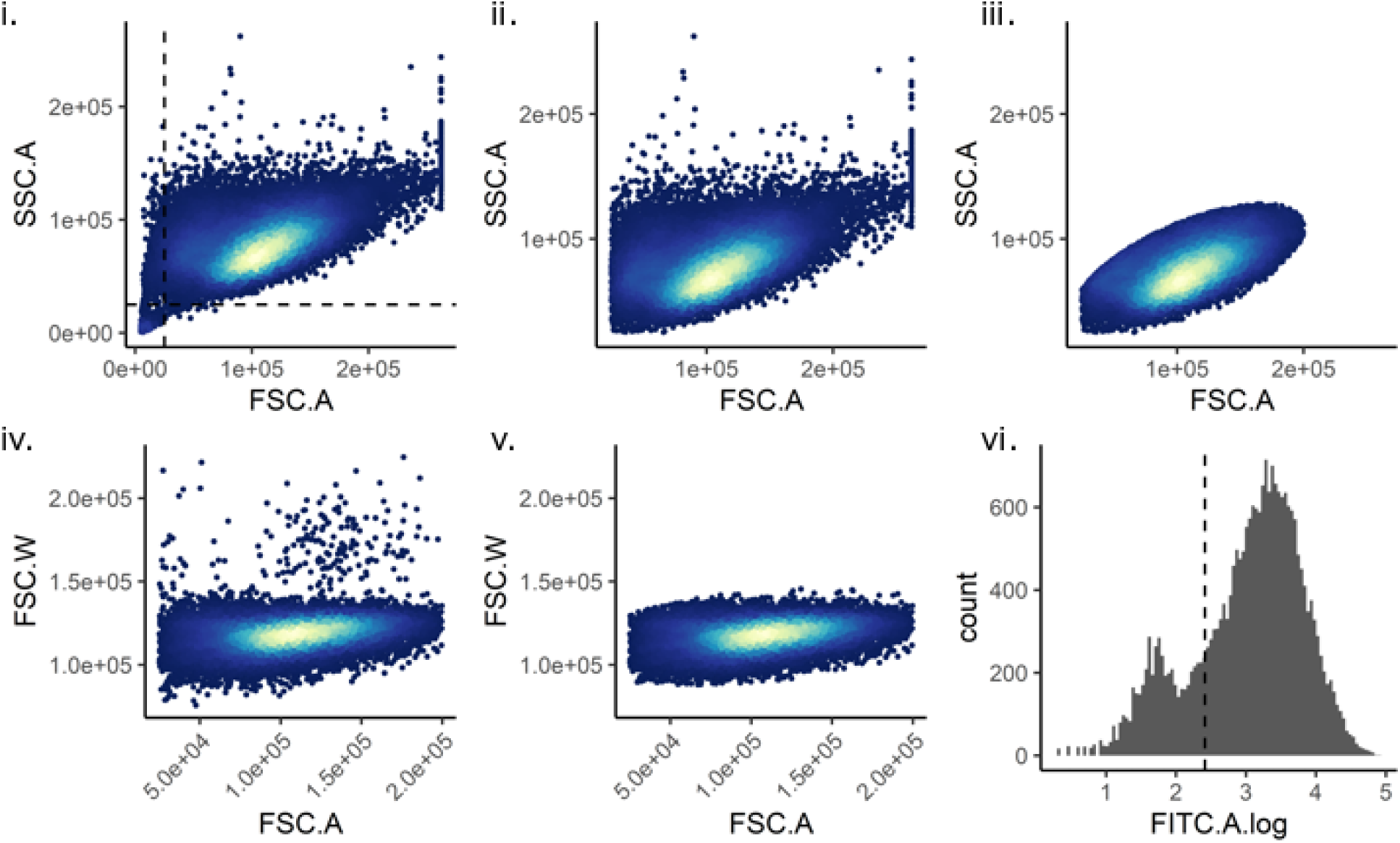

Custom code using flowCore (v1.11.20) in R (v3.6.0) was used to gate the cells as follows. (i) Debris was removed using a SSC.A vs FSC.A plot. (ii) and (iii) A Mahalanobis distance filter was used to identify live cells on a SSC.A vs FSC.A plot. (iv) and (v) A Mahalanobis distance filter was used to identify single cells on a FSC.W vs FSC.A plot. (vi) Dendra2-positive cells were identified using a FITC plot.

### Extended description of the gating scheme for Visual Cell Sorting of cells expressing the SV40 NLS library

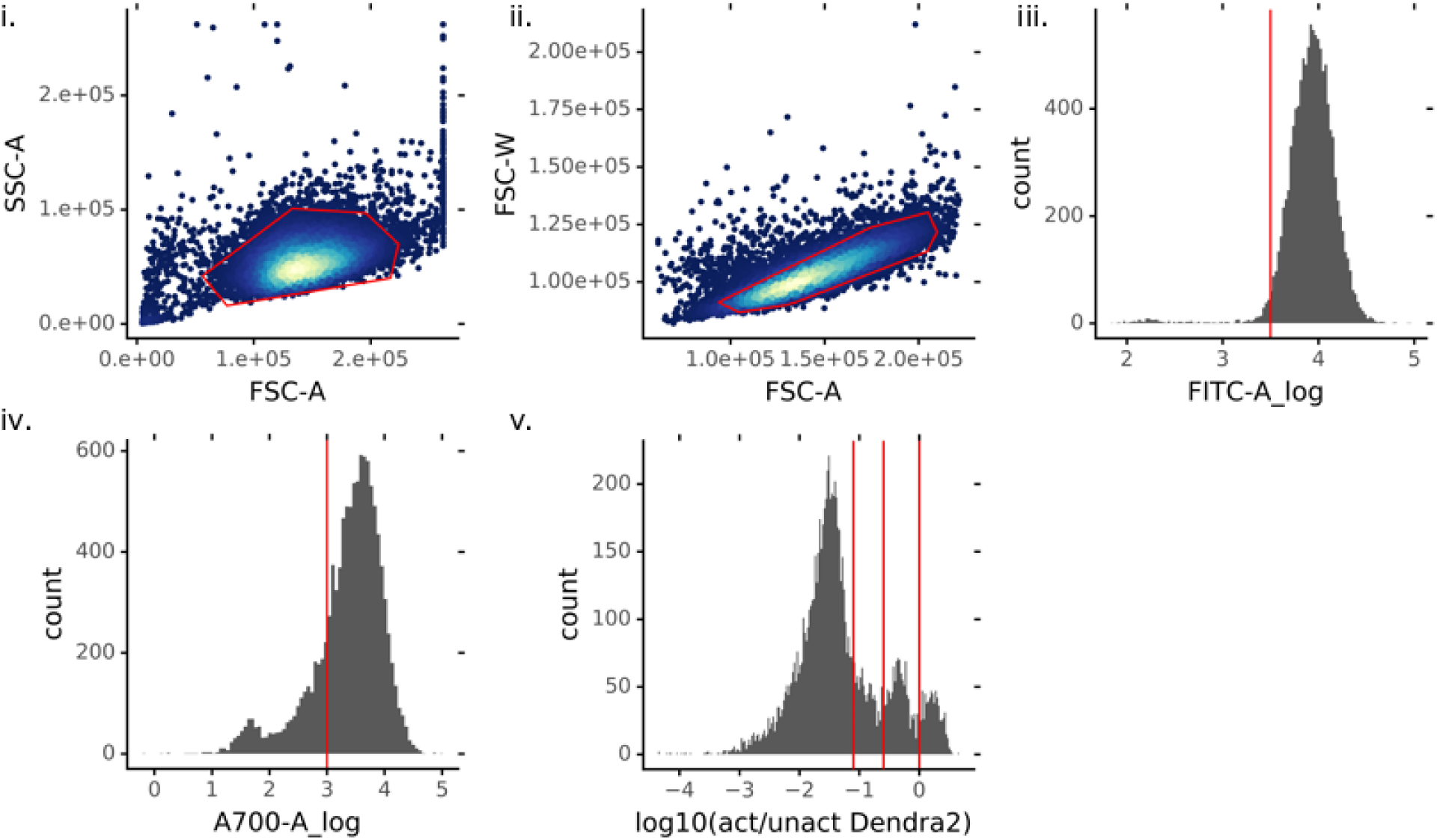

Using the BD FACSDiva software, cells were gated as follows. (i) Live cells were identified using a SSC-A vs FSC-A plot (ii) Single cells were gated using a FSC-W vs FSC-A plot. (iii) Cells expressing Dendra2 were gated using a FITC-A plot. (iv) Cells expressing miRFP were gated using an AlexaFluor 700-A plot. (v) Cells were divided into four bins according to the ratio of activated / unactivated Dendra2.

### Extended description of validation of single NLS variants

Analysis of Metamorph-calculated nucleus and cytoplasm mean intensity values was done using Python (v3.6.5). To correct for differences in background intensity between wells and replicates, each image’s miRFP background intensity was estimated using the 10^th^ percentile of image pixel intensity values and this value was subtracted from each cell’s mean nucleus and cytoplasmic miRFP intensity. Cells with no miRFP expression were removed with a gate that was determined by examination of the histogram of the mean miRFP intensity values for cells in each well. An iPython notebook file with the code used to run the analysis is available on the GitHub repository.

### Extended description of the Visual Cell Sorting on cells treated with paclitaxel

To identify morphologically-normal and lobulated cells were imaged for unactivated Dendra2 (FITC channel; 100 ms). Then, a custom nuclear segmentation pipeline that optimizes detection of nuclear blebs, herniations, and other abnormalities was employed. First, a top hat filter with a maximum object area threshold of 5,000 pixels was applied to remove large autofluorescent objects, and a 3×3 low pass filter was applied to smooth nuclear fluorescence. To find nuclei, a flatten background filter (removal of objects < 20 pixels in size), Sobel edge detection kernel, and a sharpening kernel were used before applying Metamorph’s “legacy heuristic” thresholding algorithm to create nuclear binaries. To clean the nuclear binaries, holes were filled; tunnels 1 pixel in width were filled in using a dilate function; holes were filled again; and then an erode function was used to reverse the enlarging effect of the dilate and edge detection steps. Finally, objects less than 20 pixels in size and greater than 400 pixels in size were discarded. Shape factors were computed for each remaining object. See the GitHub repository for the Metamorph journal that was used.

### Extended description of the gating scheme for Visual Cell Sorting on cells treated with paclitaxel

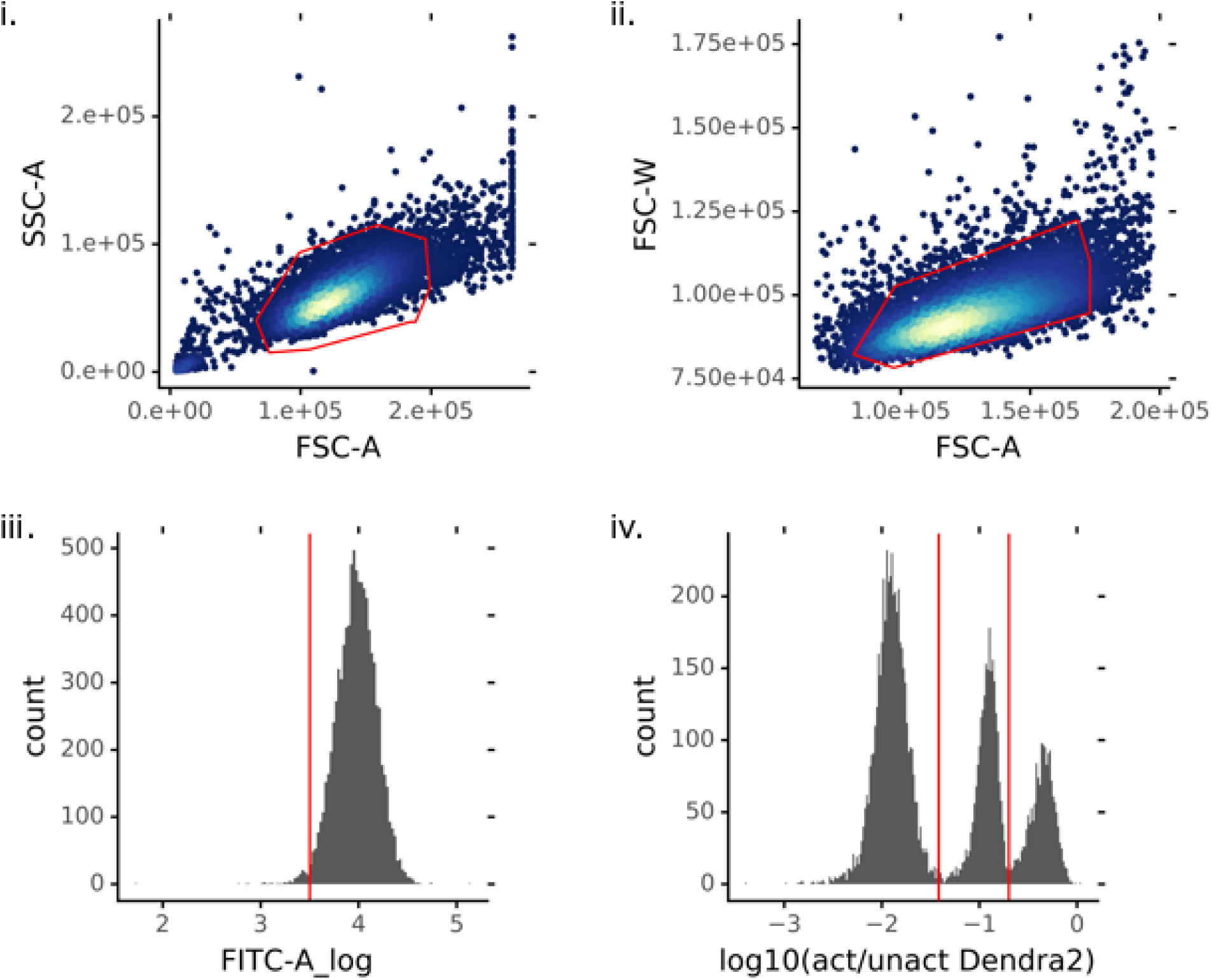

Using the BD FACSDiva software, cells were gated as follows. (i) Live cells were identified using a SSC-A vs FSC-A plot (ii) Single cells were gated using a FSC-W vs FSC-A plot. (iii) Cells expressing Dendra2 were gated using a FITC-A plot. (iv) Cells were divided into three bins (0 ms, 200 ms, and 800 ms) according to the ratio of activated / unactivated Dendra2. Cells in the 200 ms and 800 ms bins were sorted.

### Extended Description of the Differentially Expressed Genes Analysis

We noted that the Visual Cell Sorting-derived lobulated and normal single cell RNA transcriptomes appeared to be confounded by a batch effect, despite the fact that cells were derived from a single well, sorted on the same day, and processed side by side (Experiment 1; Supplementary Figure 4). Using SoupX^27^, which applies a linear PCA transformation that is determined by the RNA in empty 10X emulsion droplets, we found that cell-free RNA was responsible for this effect (Supplementary Figure 4). To confirm this hypothesis, we repeated the experiment but sorted lobulated (800 ms activated) and morphologically normal (200 ms activated cells) cells into the same bin (Experiment 2, “unseparated” population) and processed these cells in a single 10X lane. A UMAP embedding of the single cell transcriptomes derived from these unseparated cells showed a single cluster, confirming the batch effect in Experiment 1.

Although both SoupX and the mutual nearest neighbors algorithm^32^ applied to cells in Experiments 1 corrected the batch effect (Supplementary Figure 4), it is not statistically appropriate to use batch-corrected gene expression values to conduct a differentially-expressed gene test^32^. As such, we sought to use mutual nearest neighbors batch-corrected principle components to label the unseparated cells in Experiment 2 according to their similarity to the known lobulated or morphologically-normal cells in Experiment 1; and then use the raw gene expression values in Experiment 2 to conduct a differentially-expressed gene test. We noted that the first four principle components in the MNN batch correction output correlated with the visual phenotype (i.e. morphologically normal vs. lobulated) in Experiment 1. So, we performed on the cells in Experiment 1 a logistic regression to devise a score that distinguishes between morphologically normal and lobulated cells:

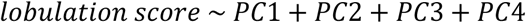

This regression model was then applied to cells in Experiment 2 (unseparated cells, single 10X lane) and the model predictions, which we called the “lobulation scores”, were extracted for each cell. Using Moncole3^55, 56^, we performed on the Experiment 2 gene expression matrix a DEG test using the lobulation scores and Seurat-computed cell cycle scores as covariates:

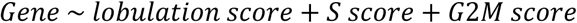

By doing the differentially expressed gene test using Experiment 2, in which lobulated and normal cells were sequenced together, we avoided any batch-related artifacts. This operation is analogous to the cluster-based analysis originally discussed by Haghverdi and colleagues^32^, but uses a principle component-derived score rather than principle component-derived clusters as cell labels.

## References

1. Boutros, M., Heigwer, F. & Laufer, C. Microscopy-Based High-Content Screening. Cell 163, 1314–1325 (2015).

2. Chen, K. H. et al. Spatially resolved, highly multiplexed RNA profiling in single cells. Science 348, aaa6090 (2015).

3. Moffitt, J. R. et al. High-throughput single-cell gene-expression profiling with multiplexed error-robust fluorescence in situ hybridization. Proc. Natl. Acad. Sci. 113, 11046–11051 (2016).

4. Emanuel, G., Moffitt, J. R. & Zhuang, X. High-throughput, image-based screening of pooled genetic-variant libraries. Nat. Methods 14, 1159–1162 (2017).

5. Wang, C., Lu, T., Emanuel, G., Babcock, H. P. & Zhuang, X. Imaging-based pooled CRISPR screening reveals regulators of lncRNA localization. Proc. Natl. Acad. Sci. U. S. A. 166, 10842–10851 (2019).

6. Eng, C.-H. L. et al. Transcriptome-scale super-resolved imaging in tissues by RNA seqFISH+. Nature (2019). doi:10.1038/s41586-019-1049-y

7. Lee, J. H. et al. Sequencing in Situ. Science 343, 1360–1363 (2014).

8. Lee, J. H. et al. Fluorescent in situ sequencing (FISSEQ) of RNA for gene expression profiling in intact cells and tissues. Nat. Protoc. 10, 442–58 (2015).

9. Feldman, D. et al. Pooled optical screens in human cells. Cell in press, 383943 (2019).

10. Binan, L. et al. Live single-cell laser tag. Nat. Commun. 7, 1–8 (2016).

11. Chien, M.-P., Werley, C. A., Farhi, S. L. & Cohen, A. E. Photostick: a method for selective isolation of target cells from culture. Chem. Sci. 6, 1701–1705 (2015).

12. Binan, L. et al. Opto-magnetic capture of individual cells based on visual phenotypes. Elife 8, 1–21 (2019).

13. Kuo, C. T. et al. Optical painting and fluorescence activated sorting of single adherent cells labelled with photoswitchable Pdots. Nat. Commun. 7, 1–11 (2016).

14. David, E. et al. Spatial reconstruction of immune niches by combining photoactivatable reporters and scRNA-seq. Science(80-.). 358, 1622–1626 (2017).

15. Ke, R. et al. In situ sequencing for RNA analysis in preserved tissue and cells. Nat. Methods 10, 1–6 (2013).

16. Feldman, D. et al. Optical Pooled Screens in Human Cells. Cell 179, 787–799.e17 (2019).

17. Chudakov, D. M., Lukyanov, S. & Lukyanov, K. A. Tracking intracellular protein movements using photoswitchable fluorescent proteins PS-CFP2 and Dendra2. Nat. Protoc. 2, 2024–2032 (2007).

18. Lin, J. rong & Hu, J. SeqNLS: nuclear localization signal prediction based on frequent pattern mining and linear motif scoring. PLoS One 8, (2013).

19. Nguyen Ba, A. N., Pogoutse, A., Provart, N. & Moses, A. M. NLStradamus: A simple Hidden Markov Model for nuclear localization signal prediction. BMC Bioinformatics 10, 1–11 (2009).

20. Kalderon, D., Roberts, B. L., Richardson, W. D. & Smith, A. E. A short amino acid sequence able to specify nuclear location. Cell 39, 499–509 (1984).

21. Rubin, A. F. et al. A statistical framework for analyzing deep mutational scanning data. Genome Biol. 18, 1–15 (2017).

22. Bloom, J. D. An Experimentally Determined Evolutionary Model Dramatically Improves Phylogenetic Fit Article Fast Track. 31, 1956–1978 (2014).

23. Thul, P. J. et al. A subcellular map of the human proteome. Science (80-.). 356, eaal3321 (2017).

24. Rowinsky, E. K., Onetto, N., Canetta, R. M. & Arbuck, S. G. Taxol: the first of the taxanes, an important new class of antitumor agents. Semin. Oncol. 19, 646–62 (1992).

25. Theodoropoulos, P. A., Polioudaki, H., Kostaki, O., Dargemont, C. & Georgatos, S. D. Taxol Affects Nuclear Lamina and Pore Complex Organization and Inhibits Import of Karyophilic Proteins into the Cell Nucleus Taxol Affects Nuclear Lamina and Pore Complex Organization and Inhibits Import of Karyophilic Proteins into the Cell Nucleus 1. 4625–4633 (1999).

26. Rohban, M. H. et al. Systematic morphological profiling of human gene and allele function via Cell Painting. Elife 1–23 (2017). doi:10.7554/eLife.24060

27. Young, M. D. & Behjati, S. SoupX removes ambient RNA contamination from droplet based single cell RNA sequenc-ing data. bioRxiv 303727 (2018). doi:10.1101/303727

28. Cao, J. et al. Comprehensive single-cell transcriptional profiling of a multicellular organism. Science (80-.). 357, 661–667 (2017).

29. McInnes, L., Healy, J. & Melville, J. UMAP: Uniform Manifold Approximation and Projection for Dimension Reduction. (2018).

30. Butler, A., Hoffman, P., Smibert, P., Papalexi, E. & Satija, R. Integrating single-cell transcriptomic data across different conditions, technologies, and species. Nat. Biotechnol. 36, 411–420 (2018).

31. Trielli, M. O., Andreassen, P. R., Lacroix, F. B. & Margolis, R. L. Differential taxol-dependent arrest of transformed and nontransformed cells in the G1 phase of the cell cycle, and specific-related mortality of transformed cells. J. Cell Biol. 135, 689–700 (1996).

32. Haghverdi, L., Lun, A. T. L., Morgan, M. D. & Marioni, J. C. Batch effects in single-cell RNA-sequencing data are corrected by matching mutual nearest neighbors. 36, (2018).

33. Gasic, I., Boswell, S. A. & Mitchison, T. J. Tubulin mRNA stability is sensitive to change in microtubule dynamics caused by multiple physiological and toxic cues. PLoS Biol. 17, e3000225 (2019).

34. Li, N., Zoubeidi, A., Beraldi, E. & Gleave, M. E. GRP78 regulates clusterin stability, retrotranslocation and mitochondrial localization under ER stress in prostate cancer. Oncogene 32, 1933–1942 (2013).

35. Cho, J. H. et al. Arginine methylation-dependent regulation of ASK1 signaling by PRMT1. Cell Death Differ. 19, 859–870 (2012).

36. Georges, E., Bonneau, A. M. & Prinos, P. RNAi-mediated knockdown of α-enolase increases the sensitivity of tumor cells to antitubulin chemotherapeutics. Int. J. Biochem. Mol. Biol. 2, 303–308 (2011).

37. Alli, E., Yang, J. M., Ford, J. M. & Hait, W. N. Reversal of stathmin-mediated resistance to paclitaxel and vinblastine in human breast carcinoma cells. Mol. Pharmacol. 71, 1233–1240 (2007).

38. Zhou, M. et al. Warburg effect in chemosensitivity: Targeting lactate dehydrogenase-A re-sensitizes Taxol-resistant cancer cells to Taxol. Mol. Cancer 9, 1–12 (2010).

39. Di Michele, M. et al. A proteomic approach to paclitaxel chemoresistance in ovarian cancer cell lines. Biochim. Biophys. Acta - Proteins Proteomics 1794, 225–236 (2009).

40. Sugimura, M. et al. Mechanisms of paclitaxel-induced apoptosis in an ovarian cancer cell line and its paclitaxel-resistant clone. Oncology 66, 53–61 (2004).

41. Liberzon, A. et al. The Molecular Signatures Database Hallmark Gene Set Collection. Cell Syst. 1, 417–425 (2015).

42. Parasido, E. et al. The Sustained Induction of c-MYC Drives Nab-Paclitaxel Resistance in Primary Pancreatic Ductal Carcinoma Cells. Mol. Cancer Res. 17, 1815–1827 (2019).

43. Shafer, A., Zhou, C., Gehrig, P. A., Boggess, J. F. & Bae-Jump, V. L. Rapamycin potentiates the effects of paclitaxel in endometrial cancer cells through inhibition of cell proliferation and induction of apoptosis. Int. J. Cancer 126, 1144–1154 (2010).

44. Shcherbakova, D. M. et al. Bright monomeric near-infrared fluorescent proteins as tags and biosensors for multiscale imaging. Nat. Commun. 7, 1–12 (2016).

45. Yang, J. et al. mBeRFP, an Improved Large Stokes Shift Red Fluorescent Protein. PLoS One 8, 6–11 (2013).

46. Ounkomol, C., Seshamani, S., Maleckar, M. M. & Collman, F. Label-free prediction of three-dimensional fluorescence images from transmitted light microscopy. (2018). doi:10.1101/289504

47. Christiansen, E. M. et al. In Silico Labeling: Predicting Fluorescent Labels in Unlabeled Images. Cell 1–12 (2018). doi:10.1016/j.cell.2018.03.040

48. Bray, M.-A. et al. Cell Painting, a high-content image-based assay for morphological profiling using multiplexed fluorescent dyes. Nat. Protoc. 11, 1757–1774 (2016).

49. Gibson, D. G. et al. Enzymatic assembly of DNA molecules up to several hundred kilobases. Nat. Methods 6, 343–5 (2009).

50. Matreyek, K. A., Stephany, J. J. & Fowler, D. M. A platform for functional assessment of large variant libraries in mammalian cells. Nucleic Acids Res. 45, (2017).

51. Matreyek, K. A., Stephany, J. J., Chiasson, M. A., Hasle, N. & Fowler, D. M. An improved platform for functional assessment of large protein libraries in mammalian cells. Nucl. Acids Res. Preprint, (2019).

52. Love, M. I., Huber, W. & Anders, S. Moderated estimation of fold change and dispersion for RNA-seq data with DESeq2. Genome Biol. 15, 1–21 (2014).

53. Zhang, J., Kobert, K., Flouri, T. & Stamatakis, A. PEAR: A fast and accurate Illumina Paired-End reAd mergeR. Bioinformatics 30, 614–620 (2014).

54. Matreyek, K. A. et al. Multiplex assessment of protein variant abundance by massively parallel sequencing. Nat. Genet. 50, 874–882 (2018).

55. Trapnell, C. et al. The dynamics and regulators of cell fate decisions are revealed by pseudotemporal ordering of single cells. Nat. Biotechnol. 32, 381–386 (2014).

56. Qiu, X. et al. Single-cell mRNA quantification and differential analysis with Census. Nat. Methods 14, 309–315 (2017).

57. Väremo, L., Nielsen, J. & Nookaew, I. Enriching the gene set analysis of genome-wide data by incorporating directionality of gene expression and combining statistical hypotheses and methods. Nucleic Acids Res. 41, 4378–4391 (2013).

58. Subramanian, A. et al. Gene set enrichment analysis: A knowledge-based approach for interpreting genome-wide expression profiles. Proc. Natl. Acad. Sci. 102, 15545–15550 (2005).

59. Ooe, A., Kato, K. & Noguchi, S. Possible involvement of CCT5, RGS3, and YKT6 genes up-regulated in p53-mutated tumors in resistance to docetaxel in human breast cancers. Breast Cancer Res. Treat. 101, 305–315 (2007).

60. Su, D. et al. Stathmin and tubulin expression and survival of ovarian cancer patients receiving platinum treatment with and without paclitaxel. Cancer 115, 2453–2463 (2009).

61. Dorman, S. N. et al. Genomic signatures for paclitaxel and gemcitabine resistance in breast cancer derived by machine learning. Mol. Oncol. 10, 85–100 (2016).

